# Minimally invasive myocardial infarction model recapitulates patient immune responses and reveals a pathogenic role for immature neutrophils

**DOI:** 10.64898/2026.03.05.709757

**Authors:** Zoe Möller-Ramon, Anna Kaltenbach, Sarah-Lena Puhl, Immanuel Kwok, Florian Sicklinger, Yvonne Jansen, Annick Ernst, Katrin Nitz, Maximilian J. Schloss, Florian Leuschner, Mark Yan Yee Chan, Christian Weber, Sabine Steffens, Johan Duchêne

## Abstract

Myocardial infarction (MI) triggers a systemic neutrophil response, yet the roles of distinct neutrophil subsets in cardiac remodeling remain unclear. Studying this requires murine models that accurately mirror human neutrophil dynamics. Here, we show that a minimally invasive intact-chest MI model is more pathophysiologically relevant than the standard open-chest approach for investigating post-MI immune responses. In the open-chest model, surgical trauma disrupts bone marrow homeostasis, releases large numbers of immature neutrophils, and masks MI-specific immune mechanisms. In contrast, the intact-chest model preserves bone marrow integrity and induces only a modest rise in circulating immature neutrophils, closely reflecting MI patient profiles. We further demonstrate that accumulation of immature neutrophils in the infarcted heart exacerbates cardiac dysfunction. Beyond neutrophils, the overall cardiac immune landscape differs markedly between both models. Collectively, our findings establish the intact-chest model as superior for studying post-MI inflammation and reveal immature neutrophils as mediators of adverse cardiac remodeling.

## Introduction

Myocardial infarction (MI), a leading cause of death and disability worldwide^1^, triggers a complex inflammatory cascade that profoundly influences cardiac repair and patient outcomes^2-6^. Neutrophils represent the predominant early-responding leukocyte population recruited to the infarcted myocardium^2-12^.

Neutrophils are produced in the bone marrow (BM) through granulopoiesis, a tightly regulated process in which myeloid precursors develop first into immature and then mature neutrophils^13-15^. Under homeostatic conditions, only fully mature neutrophils are released into circulation, while immature cells are retained in the BM. Following tissue injury, mature neutrophils are rapidly recruited and consumed at sites of inflammation^16^. To meet increased demand, the BM responds by releasing larger numbers of neutrophils, including immature cells. The presence of immature neutrophils in peripheral blood is known as "left shift" and can arise through different mechanisms, depending on the severity of the inflammatory stimulus. In moderate inflammation, stored mature neutrophils and a limited proportion of immature neutrophils are mobilized, mainly under the influence of granulocyte colony-stimulating factor (G-CSF), a well-established regulator of neutrophil release^17^. In severe inflammation, sustained cytokine signaling triggers "emergency granulopoiesis"^18-20^, a rapid and amplified neutrophil production process in which cells are generated faster than adequate maturation allows^20-22^. Consequently, immature neutrophils are prematurely released and, in the most severe cases, may even outnumber mature neutrophils in the circulation.

Despite their clinical relevance, most studies on MI have examined neutrophils as a homogeneous population without distinguishing mature and immature subsets. Recent clinical studies have reported elevated circulating immature neutrophils in MI patients^23,24^. However, it remains unknown whether MI triggers emergency granulopoiesis, whether these immature cells infiltrate the infarcted heart, and whether they represent passive bystanders or actively influence post-infarction outcomes.

To address these questions, mouse MI models that closely mirror human immune responses, particularly those involving neutrophils, are needed. The widely used open-chest MI (oc-MI) model involves surgical intervention^25-27^ that induces systemic inflammation and may disrupt BM homeostasis, potentially confounding studies of neutrophil response. Here, we compared the ischemia-driven neutrophil response in the conventional open-chest approach with a newly developed intact-chest MI (ic-MI) model^28^, benchmarking both against human MI immune responses. Our goal was to identify the model that most accurately reflects human pathology, thereby enabling rigorous investigation of immature neutrophils in post-infarction setting. Defining these mechanisms may reveal novel translational targets to improve post-MI outcomes.

## Results

### Presence of circulating immature neutrophils in AMI patients

To validate recent reports of circulating immature neutrophil subsets in the context of acute MI (AMI) in participants, we assessed neutrophil maturation status in peripheral blood samples from an independent cohort of patients hospitalized for AMI (Fig. 1A-B and Extended Data Fig. 1). Flow cytometric analysis performed on peripheral venous blood obtained 24-48h from the time of symptom onset revealed a marked increase in circulating immature CD10⁻ neutrophils (iNeu), comprising approximately 10% of the total neutrophil pool compared to roughly 1% in healthy donors (Fig. 1C-D). This expansion was further evidenced by the increased proportion of CD10⁻ immature neutrophils among total CD45⁺ leukocytes (Fig. 1E) and elevated absolute counts of CD10⁻ immature neutrophils in blood of AMI patients (Fig. 1F). In contrast, the fraction of CD10^+^ mature neutrophils (mNeu) among CD45⁺ leukocytes remained unchanged (Fig. 1E), and their absolute counts were modestly elevated in these AMI patients (Fig. 1F). These findings confirm the presence of circulating immature neutrophils in AMI patients and suggest their release to be triggered by the infarction event.

**Figure 1.**
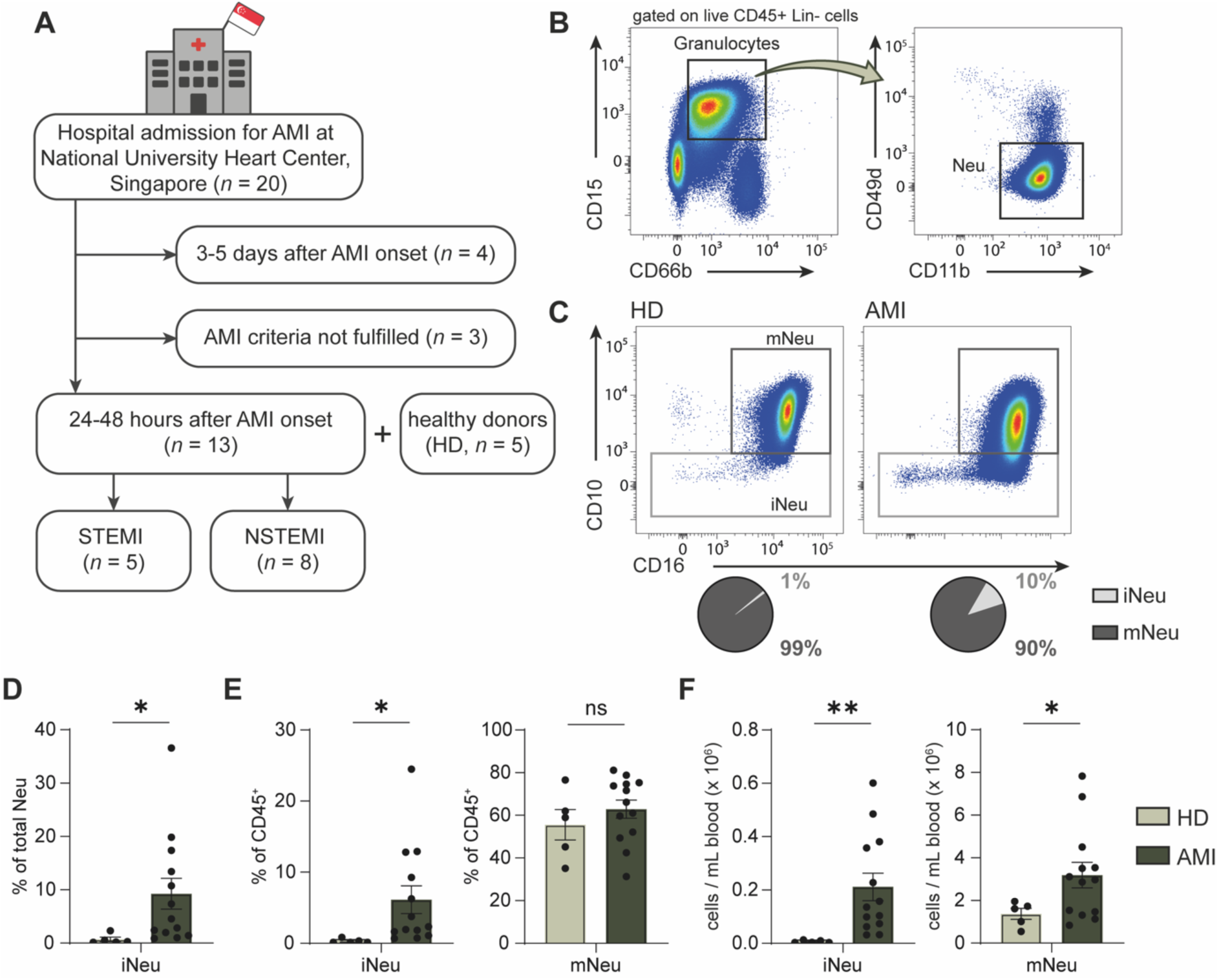
Presence of circulating immature neutrophils in AMI patients. (**A**) Schematic overview of the human study. Patients with acute myocardial infarction (AMI) were hospitalized at the National University Heart Center, Singapore (*n* = 20). Blood samples collected within 48h post-MI from patients (AMI, *n* = 13) and from healthy donors (HD, *n* = 5) were analyzed by flow cytometry. (**B**) Representative flow cytometry analysis to identify circulating neutrophils. Neutrophils were gated on the basis of CD45, CD15, CD66b and CD11b expression. Full gating strategy is available in Extended Data Fig. 1. (**C**) Representative flow cytometry analysis to distinguish neutrophil subsets based on CD10 expression in blood from donors (HD) and AMI patients. Pie charts represent the relative proportions of mature (CD10^+^) and immature (CD10^-^) subsets within total neutrophils. (**D**) Proportion of immature neutrophils within total circulating neutrophils. (**E**) Frequency of immature (left) and mature (right) neutrophils within CD45^+^ white blood cells. (**F**) Absolute counts of immature (left) and mature (right) neutrophils. (**D-F**) Data are shown as mean ± SEM. Welch’s *t*-test. \**p* < 0.05 and \*\**p* < 0.01. HD: *n* = 5; AMI: *n* = 13. ns, non-significant; STEMI, ST-elevated myocardial infarction; NSTEMI, non-ST-elevated myocardial infarction; Neu, neutrophils; mNeu, mature neutrophils; iNeu, immature neutrophils; Lin, Lineage (CD3, CD19, C56, CD14).

### Open-chest surgery alters bone marrow homeostasis

To test whether MI truly induces left shift and to explore the underlying mechanisms, we next employed mouse models. First, we used the widely established oc-MI model (Fig. 2A) and examined whether it recapitulates human pathology by assessing the presence of circulating immature neutrophils following left anterior descending (LAD) artery ligation (Extended Data Fig. 2A). We observed a profound increase in circulating CD101^-^ immature neutrophils, which constituted up to 80% of total neutrophils 24h post-MI, compared to 5% in steady state (Fig. 2B-C). This was associated with an elevation in CD101^-^ immature and a corresponding reduction in CD101^+^ mature neutrophil numbers (Fig. 2D). Strikingly, sham-operated mice (oc-SO) exhibited a comparable response to oc-infarcted mice, with a marked rise in CD101⁻ immature neutrophils and a near-complete loss of CD101⁺ mature neutrophils in circulation (Fig. 2B–D).

**Figure 2.**
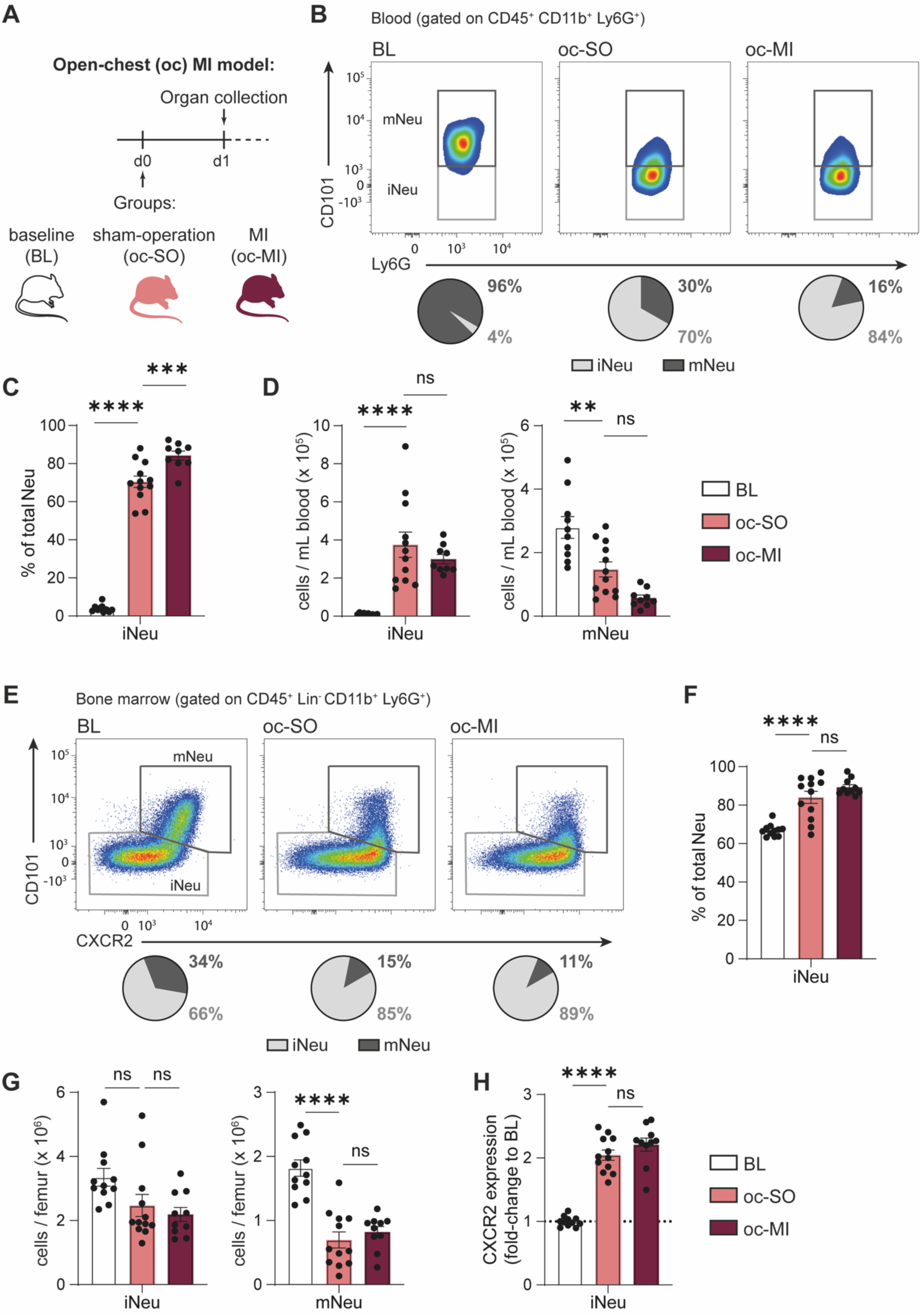
Surgery in open-chest MI model alters BM homeostasis and induces a marked increase in circulating immature neutrophils. (**A**) Experimental design in the open-chest (oc) MI model: Mice were untreated as baseline controls (BL, white), sham-operated (SO, light red), or subjected to LAD ligation (MI, dark red). Blood and bone marrow (BM) were collected 24h post-intervention for flow cytometric analysis. (**B**) Representative flow cytometry analysis to identify neutrophil subsets in blood. CD101^+^ neutrophils were classified as mature and CD101^-^ as immature. Pie charts indicate the average proportion of each subset within total circulating neutrophils. (**C-D)** Frequency of immature neutrophils within total blood neutrophils (**C**) and absolute counts of circulating immature (left) and mature (right) neutrophils (**D**). (**E**) Representative flow cytometry analysis to identify neutrophil subsets in BM. CD101^+^ neutrophils were considered mature, CD101^-^immature. Pie charts indicate the average proportion of each subset within total BM neutrophils. (**F-G**) Frequency of immature neutrophils within total BM neutrophils (**F**) and absolute counts of BM immature (left) and mature (right) neutrophils (**G**). (**H**) CXCR2 expression on immature neutrophils shown as fold-change relative to BL controls. (**B-D** and **F-H**) Data are shown as mean ± SEM. One-way ANOVA with Bonferroni *post hoc* test, \*\**p* < 0.01, \*\*\**p* < 0.001, and \*\*\*\**p* < 0.0001. BL *n* = 10, oc-SO *n* = 12 and *n* = 9 (**B-D**), BL *n* = 11, oc-MI *n* = 12, and oc-MI *n* = 10 (**F-H**). ns, non-significant; Neu, neutrophils; mNeu, mature neutrophils; iNeu, immature neutrophils.

To investigate the origin of this neutrophil shift, we next examined the BM compartment (Fig. 2E, Extended Data Fig. 2B). In both oc-MI and oc-SO groups, CD101⁺ mature neutrophils were virtually depleted at 24h post-intervention and replaced by CD101⁻ immature neutrophils (Fig. 2F-G). Kinetic analysis revealed that all mature CD101⁺ neutrophils were rapidly mobilized into circulation as early as 4h post-intervention, leaving immature neutrophils as the predominant population in the BM (Extended Data Fig. 3A-B). This abrupt change explains the overwhelming presence of immature neutrophils in both BM and peripheral blood at 24h. BM composition and blood neutrophil distribution gradually returned to baseline within seven days (Extended Data Fig. 3A-D). Additionally, CXCR2 expression, normally restricted to mature neutrophils in steady state, was significantly upregulated on CD101⁻ immature neutrophils in the BM of both oc-MI and oc-SO mice (Fig. 2H), indicating that surgical intervention alters the phenotype of immature neutrophils.

Together, these findings demonstrate that open-chest surgery alone (i.e., thoracotomy) triggers a severe immune response, disrupting granulopoiesis and causing a systemic shift in neutrophil populations. This dramatic alteration of BM homeostasis, affecting both composition and phenotype, significantly confounds the interpretation of immune responses in the oc-MI model, thereby limiting its utility for studying MI-specific immune mechanisms.

### Intact-chest MI model recapitulates human-like neutrophil profile

To minimize confounding factors introduced by the surgical procedure, we next employed the recently developed intact-chest MI (ic-MI) model^28^, which induces LAD artery occlusion through minimally invasive coagulation (Fig. 3A). While mature neutrophil counts remained unchanged, we found a modest but consistent increase in the percentage and number of immature neutrophils in the circulation 24h post-MI (Fig. 3B-D), closely mirroring the response observed in human patients (Fig. 1). Importantly, this increase was absent in the respective sham-operated mice (ic-SO), allowing for a clearer distinction between MI-specific and procedure-induced effects.

**Figure 3.**
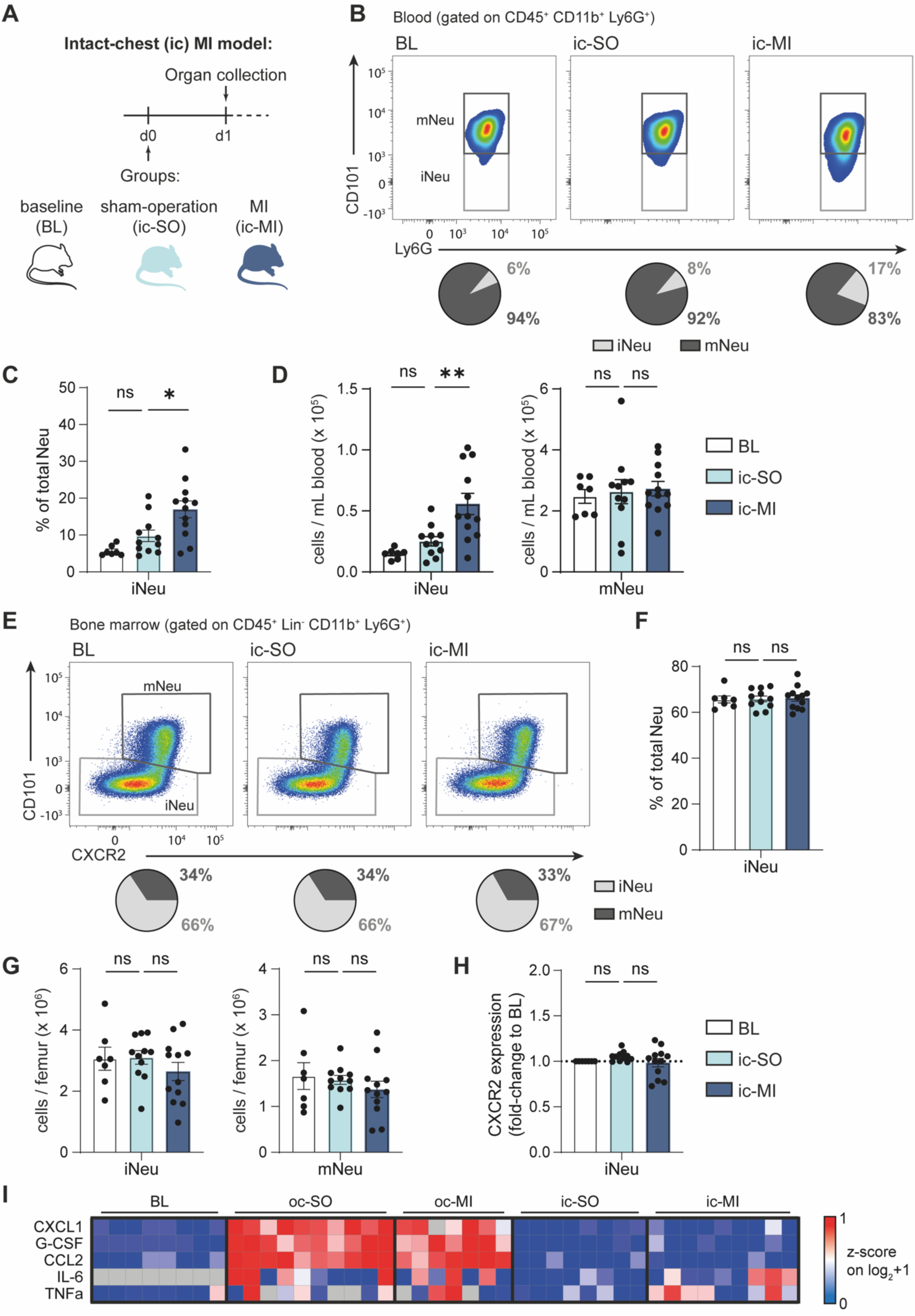
Intact-chest MI induces a moderate increase of circulating immature neutrophils. (**A**) Experimental design of the intact-chest (cc) MI model. Mice were untreated as baseline controls (BL, white), sham-operated (SO, light blue), or subjected to LAD coagulation (MI, dark blue). Blood and bone marrow (BM) were collected 24h post-intervention for flow cytometric analysis. (**B**) Representative flow cytometry analysis to identify neutrophil subsets in blood. Pie charts indicate the average proportion of CD101^+^ mature and CD101^-^ immature subsets within total circulating neutrophils. (**C-D**) Frequency of immature neutrophils within total blood neutrophils (**C**) and absolute counts of circulating immature (left) and mature (right) neutrophils (**D**). (**E**) Representative flow cytometry analysis to identify neutrophil subsets in BM. Pie charts indicate the average proportion of CD101^+^ mature and CD101^-^ immature subsets within total BM neutrophils. (**F-G**) Frequency of immature neutrophils within total BM neutrophils (**F**) and absolute counts of BM immature (left) and mature (right) neutrophils (**G**). CXCR2 expression on immature neutrophils shown as fold-change relative to BL controls. Plasma levels of CXCL1, G-CSF, CCL2, IL-6, and TNFα. Analyte concentrations were quantified by multiplex immunoassay, log₂(+1) transformed, and visualized as row z-scores. Grey boxes indicate values below the detection limit. (**B-D** and **F-H**). Data are shown as mean ± SEM. One-way ANOVA with Bonferroni *post hoc* test, \**p* < 0.05 and \*\**p* < 0.01. BL *n* = 7, ic-SO *n* = 11, and ic-MI *n* = 12. ns, non-significant; BL, baseline; Neu, neutrophils; mNeu, mature neutrophils; iNeu, immature neutrophils.

In the BM, the percentage and absolute numbers of mature and immature neutrophils were comparable between ic-MI and ic-SO 24h post-intervention and baseline (BL) mice (Fig. 3E-G). Additionally, CXCR2 expression on immature neutrophils remained unchanged after sham (ic-SO) and MI (ic-MI) procedures (Fig. 3H). Thus, by avoiding surgical confounding effects, the intact-chest model reveals that MI does not substantially alter BM neutrophil homeostasis.

To further dissect the divergent responses between the open-chest and intact-chest models, we quantified circulating cytokines and chemokines 24h post-intervention using a multiplex immunoassay (Fig. 3I). Sham surgery alone in the oc-model (oc-SO) was sufficient to induce a robust upregulation of G-CSF, the key cytokine regulating neutrophil mobilization and production, and CXCL1, a chemokine involved in neutrophil trafficking, in comparison to BL controls. Similar levels of both mediators were detected in oc-infarcted (oc-MI) mice. In contrast, these mediators remained unchanged in both ic-SO and ic-MI mice, relative to BL. In addition, the chemokine CCL2 was also elevated in open-chest groups (oc-SO and oc-MI) but remained at baseline levels in the intact-chest model (ic-SO and ic-MI). Furthermore, inflammatory cytokines IL-6 and TNFα were increased in all groups except in ic-SO mice, further underscoring the dominant contribution of the surgical trauma to the systemic inflammatory response observed in the open-chest model.

Collectively, these results show that the intact-chest MI model does not provoke systemic inflammation, preserves BM integrity, and more accurately mirrors the moderate increase of circulating immature neutrophils observed in MI patients. By avoiding surgical confounding effects, the ic-MI model thus provides a powerful tool for investigating MI-specific immune responses, particularly those involving neutrophil subsets.

### MI triggers acute and rapid mobilization of immature neutrophils

Next, we investigated the origin of immature neutrophils following MI in the intact-chest model. Although we detected the presence of immature neutrophils in circulation 24h post-MI (Fig. 3B-D), neither their egress from the BM (Fig. 3E-G) nor upregulation of neutrophil-attracting molecules was observed at this time point (Fig. 3I). We therefore hypothesized that MI may induce the mobilization of immature neutrophils at earlier stages.

Time-course analysis confirmed this hypothesis as plasma levels of G-CSF, CXCL1 and IL-6 measured by ELISA peaked at 4h after ic-MI compared to ic-SO controls and returned to baseline levels within 24 hours (Fig. 4A-C). This early elevation of neutrophil chemoattractant and mobilizer molecules was followed by a significant reduction of mature and immature neutrophils in the BM at 12h post-MI (Fig. 4D–F and Extended Data Fig. 4A). Concurrently, numbers of circulating neutrophils were higher in the ic-MI group than in ic-SO controls at 12h post-MI (Fig. 4G), with an increased proportion and absolute number of immature neutrophils in circulation (Fig. 4H and I and Extended Data Fig. 4B and C).

**Figure 4.**
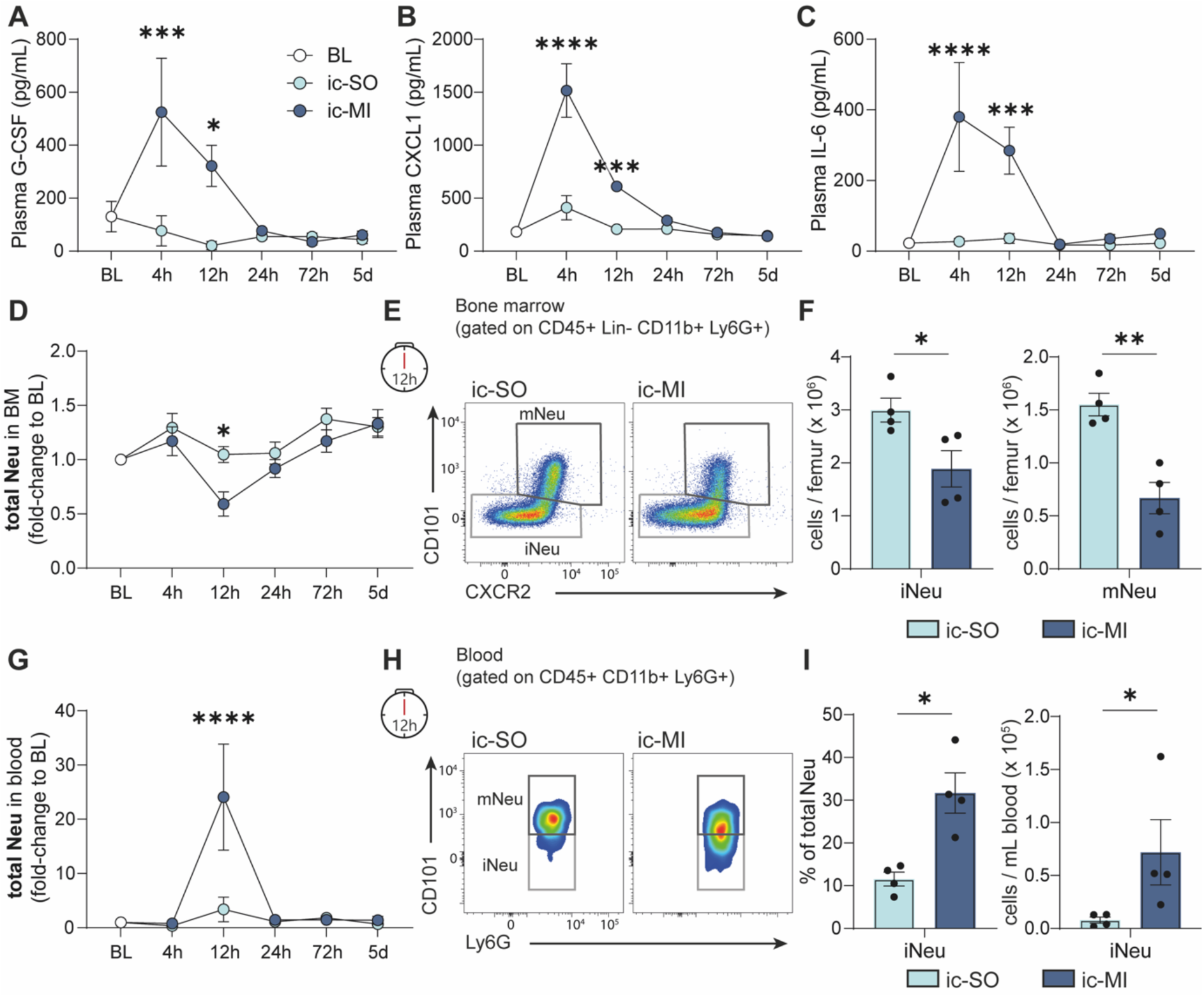
Dynamics of neutrophil response in the intact-chest MI model. In the intact-chest (cc) model, mice were sham-operated (ic-SO, light blue) or subjected to myocardial infarction (ic-MI, dark blue). Samples were collected at the indicated time points. (**A-C**) Time course of plasma levels of G-CSF, CXCL1 and IL-6 in ic-SO and ic-MI mice. Analyte concentrations were measured by ELISA. (**D**) Time course of bone marrow (BM) neutrophils in ic-SO and ic-MI mice normalized to counts of corresponding BL controls. (**E**) Representative flow cytometry analysis to identify BM neutrophil subsets 12h post-intervention. CD101^+^ neutrophils were considered mature, CD101^-^ neutrophils immature. (**F**) Quantification of immature (left) and mature (right) BM neutrophils 12h post-intervention. (**G**) Time-course of circulating neutrophils in ic-SO and ic-MI mice normalized to counts of corresponding BL controls. (**H**) Representative flow cytometry analysis to identify blood neutrophil subsets 12h post-intervention. CD101^+^ neutrophils were considered mature, CD101^-^ neutrophils immature. (**I**) Fraction of total neutrophils (left) and counts (right) of immature neutrophils in blood 12h post-intervention. (**A-D** and **G**) Data are shown as mean ± SEM. Two-way ANOVA with Šidák *post hoc* test; \**p* < 0.05, *** *p* < 0.001, and \*\*\*\**p* < 0.0001. (**A-C**) *n* = 3-6 per time point per group, except ic-SO 24h *n* = 7 and ic-MI 24h *n* = 11. (**D** and **G**) *n* = 4-7 per time point per group, except ic-SO 24h *n* = 11 and ic-MI 24h *n* = 12. (**F** and **I**) Data are shown as mean ± SEM. Mann-Whitney or Welch’s *t*-test. \**p* < 0.05 and \*\**p* < 0.01. ic-SO *n* = 4 and ic-MI *n* = 4. ns, non-significant; BL, baseline; Neu, neutrophils; mNeu, mature neutrophils; iNeu, immature neutrophils.

In conclusion, these findings indicate that MI triggers a rapid, acute, and modest mobilization of immature neutrophils from the BM. The intact-chest model thus enables temporal resolution of neutrophil dynamics following MI and demonstrates that MI induces a left shift through mobilization rather than increased granulopoiesis.

### Infiltration of immature neutrophils into the infarcted heart in ic-MI model

Next, we aimed to characterize spatial neutrophil dynamics following MI. To this end, we assessed neutrophil distribution across different organs 24h post-MI in the intact-chest model using flow cytometry. As expected, we found that MI induced neutrophil accumulation exclusively in thoracic organs, with the heart representing the primary site, followed by mediastinal lymph nodes and the lung (Fig. 5A).

**Figure 5.**
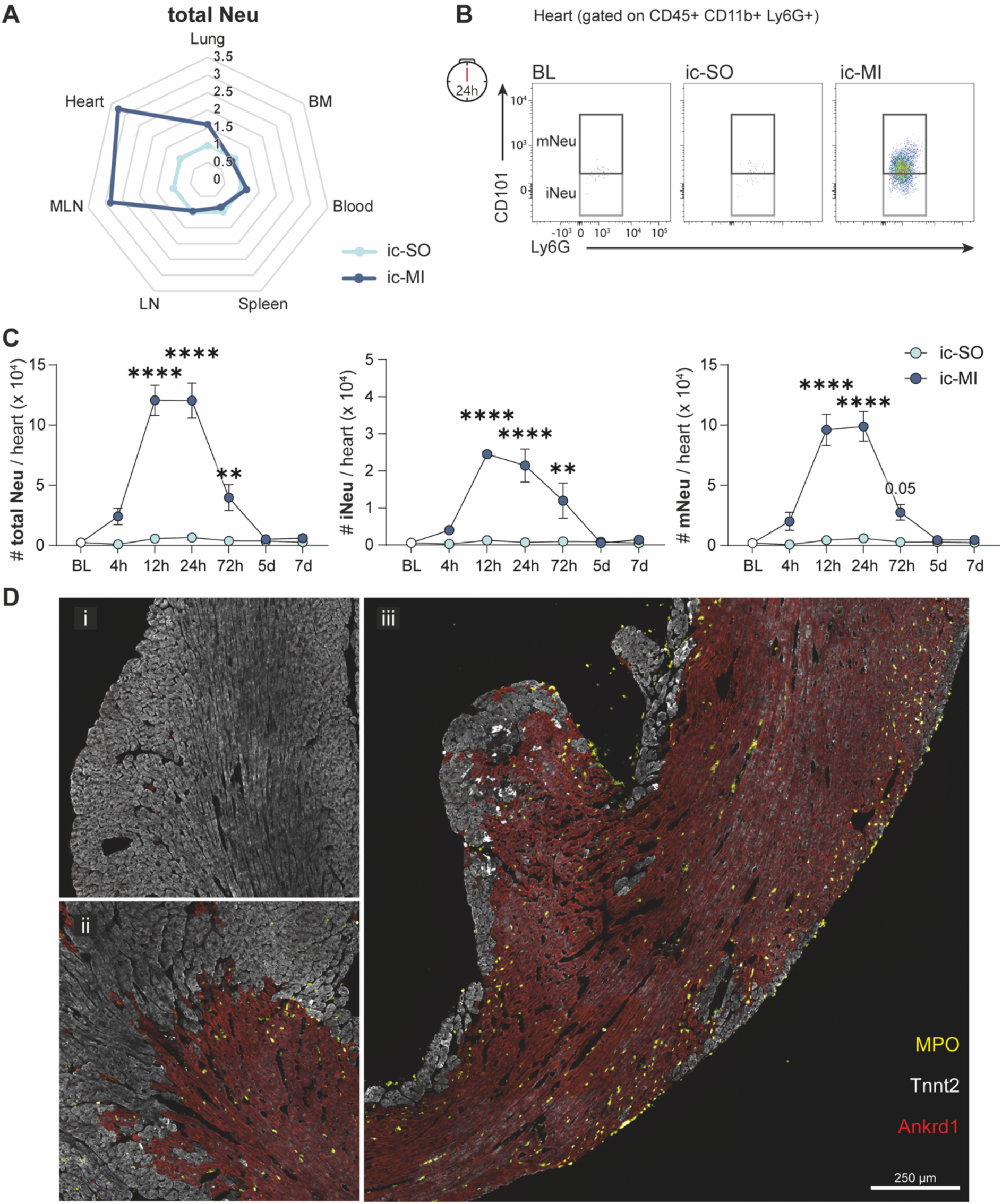
Infiltration of immature neutrophils in the infarcted heart in the intact-chest model. (**A**) Radar plot showing neutrophils distribution in organs 24h post-intervention in ic-SO and ic-MI mice. Values for ic-MI (dark blue) are expressed as fold-change relative to ic-SO (light blue). Neutrophils were measured by flow cytometry. *n* = 4-12 per organ per group. (**B**) Representative flow cytometry plots of neutrophil subsets in heart 24h post-intervention in ic-SO and ic-MI mice. (**C**) Time course of cardiac neutrophil infiltration. Absolute counts of total (left), immature (middle) and mature (right) neutrophils in the heart of ic-SO (light blue) and ic-MI (dark blue) mice at the indicated time points post-intervention. Data are shown as mean ± SEM. Two-way ANOVA between ic-SO and ic-MI groups for each time-point corrected with Šidák *post hoc* test; \*\**p* < 0.01 and \*\*\*\**p* < 0.0001. *n* = 3-5 per time point per group except ic-MI 24h *n* = 8. (**D**) Representative SeqIF images of mouse heart cross-sections 24h post ic-MI, stained for neutrophils (MPO), healthy (Tnnt2) and stressed cardiomyocytes (Ankrd1). Insets show (i) remote, (ii) border and (iii) infarct zone. Scale bar: 250 µm. ns, non-significant; BL, baseline mice; ic-SO, intact-chest sham-operation; ic-MI, intact-chest myocardial infarction; Neu, neutrophils; mNeu, mature neutrophils; iNeu, immature neutrophils; BM, bone marrow, LN, peripheral lymph nodes; MLN, mediastinal lymph nodes.

We then asked whether immature neutrophils infiltrate the infarcted myocardium. We analyzed neutrophil subsets and found that both mature and immature neutrophils infiltrated the infarcted heart (Fig. 5B and Extended Data Fig. 5). Infiltration of both populations peaked between 12h and 24h post-MI and returned to baseline levels by day 5 (Fig. 5C). Consistent with systemic observations, the ratio of immature to mature neutrophils in the heart mirrored blood dynamics, with mature neutrophils approximately three-fold more abundant than immature neutrophils in the infarcted myocardium (Fig. 4G–I and Fig. 5C).

We next investigated the spatial distribution of neutrophils in the infarcted heart using the spatiotemporal MI map generated by antibody-based Sequential Immunofluorescence (SeqIF; Lunaphore COMET)^29^. We found that 24h post-MI, MPO^+^ neutrophils were absent from remote areas (Fig. 5D inset i), sparse in border zones (Fig. 5D inset ii) and enriched in the infarct core (Fig. 5D inset iii). Interestingly, within the infarcted region, neutrophils were mainly localized beneath the endocardium and adjacent to the epicardium, suggesting their infiltration through two distinct entry routes (Fig. 5D inset iii).

Together, flow cytometric and immunofluorescence analyses confirm that mature but also immature neutrophils infiltrate the infarcted myocardium and reach the infarct area.

### A model to increase immature neutrophils recruitment to the infarcted heart

We next aimed to investigate the contribution of immature neutrophils to cardiac remodeling. However, this remains challenging as no specific tool currently exists to selectively inhibit their mobilization or recruitment. To address this limitation, we sought instead to enhance their recruitment to the heart. Given that G-CSF promotes BM egress of immature neutrophils^15,17^, we hypothesized that G-CSF treatment immediately after MI onset would further increase their numbers in the circulation and subsequently in the infarcted myocardium (Fig. 6A). Indeed, G-CSF-treated mice showed a marked increase in circulating CD101⁻ immature neutrophils 24h after MI, while mature neutrophil levels remained unchanged (Fig. 6B–C). This was accompanied by greater accumulation of CD101^-^ immature neutrophils in the infarcted heart compared with PBS-treated controls (Fig. 6D–E). Analysis of additional maturation markers confirmed the immature phenotype of these infiltrated CD101^-^ neutrophils, as they expressed lower levels of CD11b, CD62L and SiglecF than CD101^+^ mature cells (Fig. 6F).

**Figure 6.**
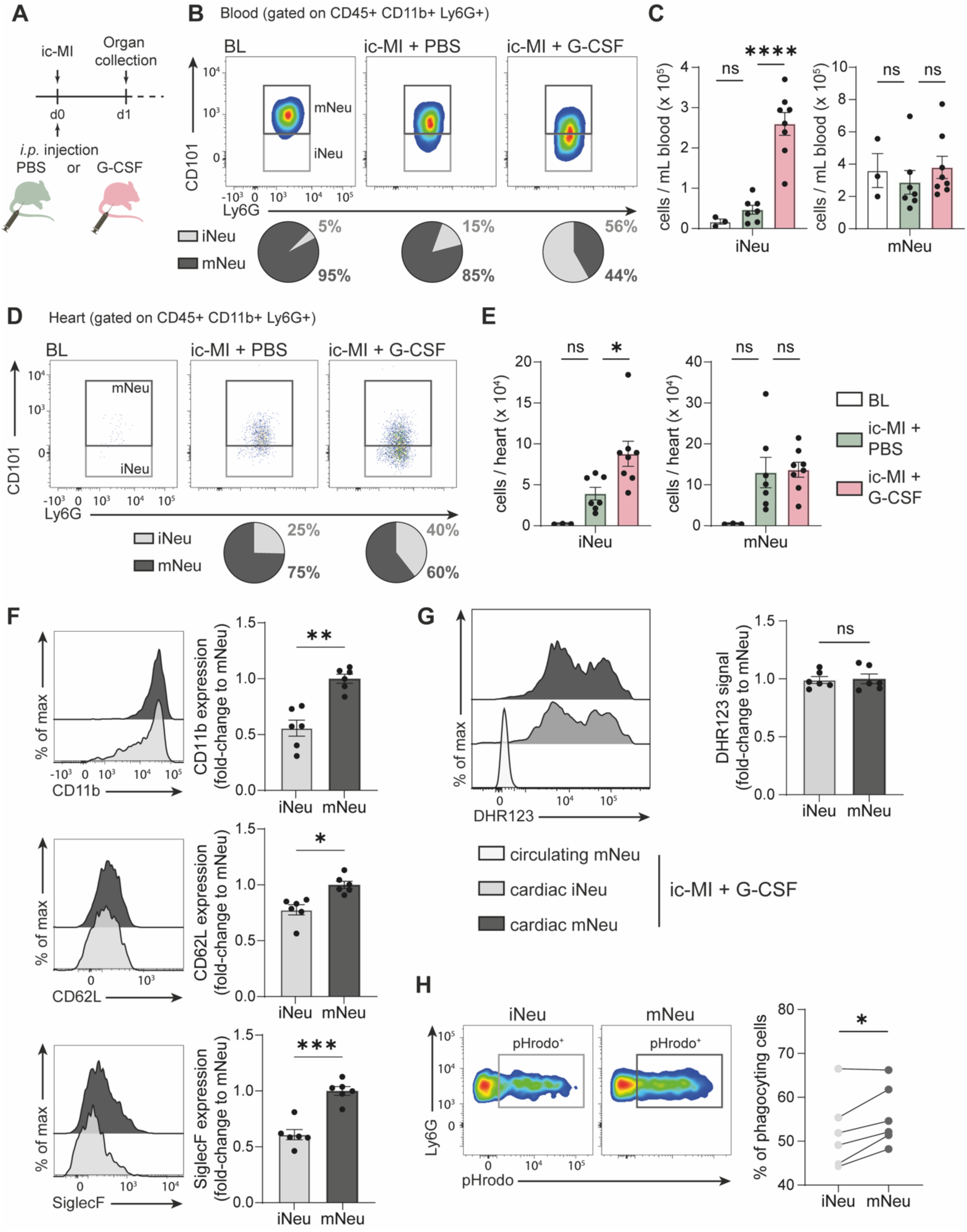
G-CSF promotes immature neutrophils infiltration into the infarcted heart. (A) Schematic of the experiment design. Mice were subjected to ic-MI and injected with either PBS (green) or G-CSF (red). Blood and hearts were collected 24h post-MI. (**B**) Representative flow cytometry analysis to identify circulating neutrophil subsets. Pie charts indicate the average proportion of CD101^+^ mature and CD101^-^ immature subsets within total circulating neutrophils. (**C**) Absolute counts of circulating immature (left) and mature (right) neutrophils from ic-MI mice treated with PBS (*n* = 7) or G-CSF (*n* = 8). (**D**) Representative flow cytometry analysis of neutrophil subsets in the heart. Pie charts indicate the average proportion of CD101^+^ mature and CD101^-^ immature subsets within total cardiac neutrophils. (**E**) Absolute counts of immature (left) and mature (right) neutrophils in infarcted hearts from ic-MI mice treated with PBS (*n* = 7) or G-CSF (*n* = 8). (**F**) Expression of CD11b, CD62L, and SiglecF on cardiac immature (light grey) and mature (dark grey) neutrophils 24h after ic-MI in G-CSF-treated mice. Representative flow cytometry histograms (left) and quantitative analysis shown as fold-change relative to mNeu (right). (**G**) ROS production by cardiac immature (light grey) and mature (dark grey) neutrophils 24h after ic-MI in G-CSF-treated mice. Representative flow cytometry histograms of DHR123 fluorescence (left) and quantitative analysis shown as fold-change relative to mNeu (right). Circulating neutrophils (white) were used as controls. (**H**) Phagocytosis capacity of cardiac immature and mature neutrophils 24h after ic-MI in G-CSF-treated mice. Representative flow cytometry histograms of pHrodo particle uptake (left) and quantitative analysis, expressed as the percentage of pHrodo-positive cells (right). (**C** and **E**) Data are shown as mean ± SEM. One-way ANOVA with Bonferroni *post hoc* test; \**p* < 0.05 and \*\*\*\**p* < 0.0001. BL *n* = 3, ic-MI+PBS *n* = 7, ic-MI+G-CSF *n* = 8. (**F**, **G** and **H**) Data are shown as mean ± SEM. Paired *t*-test with Holm-Šidák *post hoc* test; \**p* < 0.05 and \*\**p* < 0.01. Cardiac immature and mature from ic-MI+G-CSF with each *n* = 6. ns, non-significant. BL, baseline mice; ic-MI, intact-chest myocardial infarction; Neu, neutrophils; mNeu, mature neutrophils; iNeu, immature neutrophils.

To assess the functional competence of CD101^-^ immature neutrophils after cardiac infiltration, we compared their effector functions to those of CD101^+^ mature neutrophils. We first measured their ability to generate reactive oxygen species (ROS) using the cell-permeable probe dihydrorhodamine 123 (DHR123). Both mature and immature subsets isolated from the infarcted heart displayed a comparable oxidative burst that largely exceeded that of circulating mature neutrophils (Fig. 6G). Next, phagocytic capacity was evaluated using pHrodo BioParticles, which fluoresce upon internalization and phagosomal acidification. Immature neutrophils were able to internalize targets efficiently, although their phagocytic activity was modestly lower compared to mature neutrophils (Fig. 6H). Altogether, these results indicate that infiltrated immature neutrophils, despite being less differentiated, possess functional myeloid effector activities (Fig. 6G-H).

In summary, G-CSF treatment in the ic-MI model selectively enhanced the recruitment of functionally competent immature neutrophils into the infarcted heart, providing a valuable approach to investigate their role in post-infarct cardiac repair.

### High immature neutrophil counts correlate with worsened cardiac performance and adverse myocardial remodeling

To further clarify the impact of immature neutrophils on post-MI remodeling, we assessed cardiac outcomes 28 days after ic-MI in G-CSF-treated mice (Fig. 7A). Echocardiographic analysis revealed a more pronounced reduction in left ventricular ejection fraction (LVEF) and a less negative global longitudinal strain (GLS) compared to PBS-treated controls (Fig. 7B-D), indicating aggravated systolic dysfunction. Both left ventricular end-diastolic volume (LVEDV) and left ventricular end-systolic volume (LVESV) were increased in the G-CSF–treated group compared with the PBS-treated group (LVEDV: +29.7 µL, Hedges’ g = 1.18, *p* = 0.14; LVESV: +27.8 µL, Hedges’ g = 1.35, *p* = 0.10) (Fig. 7E). Although these differences did not reach conventional statistical significance (*p* > 0.05), the large observed effect sizes (Hedges’ g > 1) suggest a biologically substantial LV dilatation. Additionally, G-CSF-treated animals displayed an elevated heart weight to tibia length (HW/TL) ratio (Fig. 7F) supporting persistent inflammatory signaling and ongoing remodeling.

**Figure 7.**
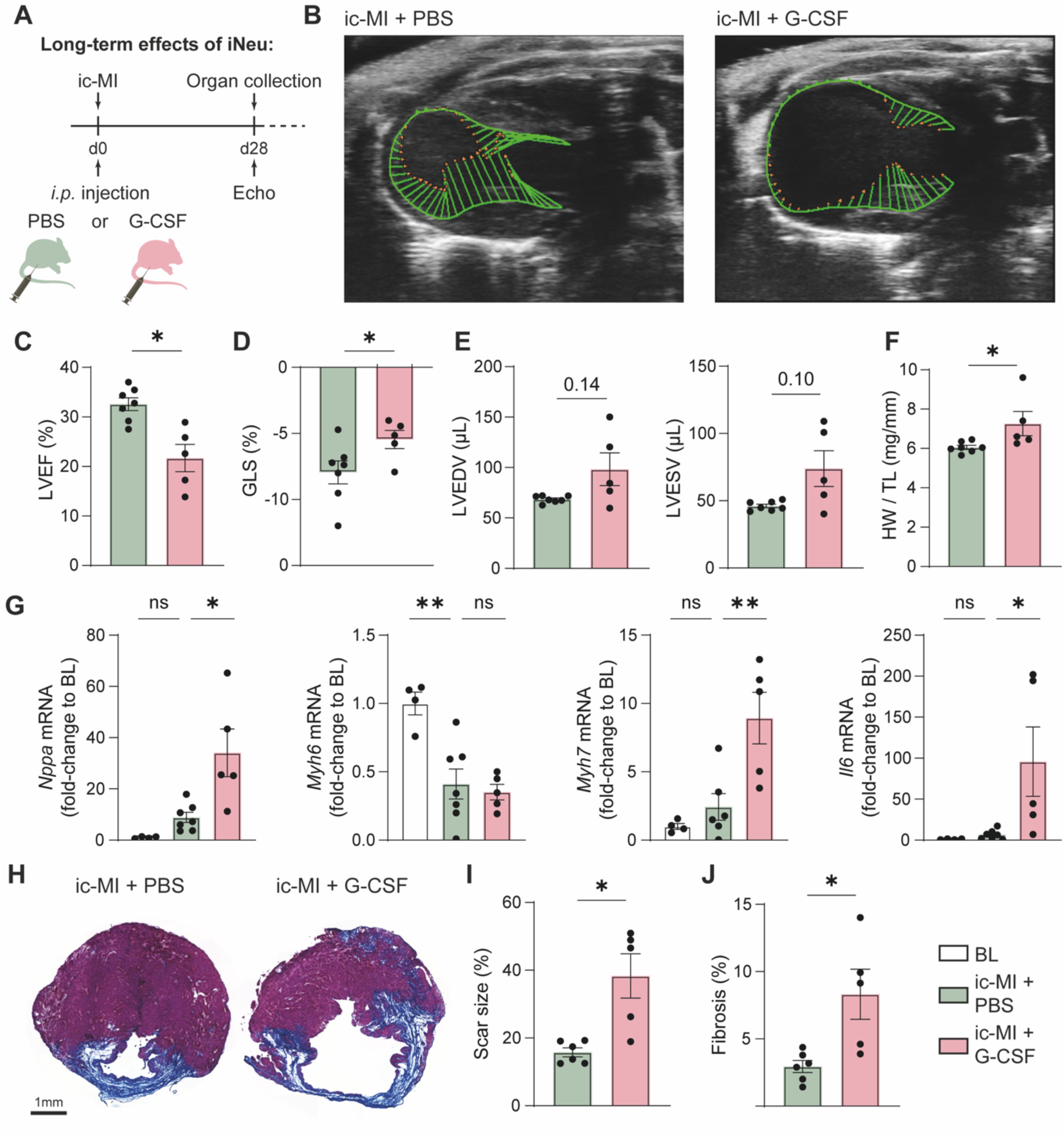
Cardiac immature neutrophil accumulation drives adverse myocardial remodeling. (A) Schematic of the experiment design. Mice were subjected to ic-MI and injected with either PBS (green) or G-CSF (red). Echocardiography and organ collection were performed 28 days post-MI. (**B-E**) Echocardiographic assessment of cardiac function in ic-MI mice treated with PBS (*n* = 7) or G-CSF (*n* = 5). (**B**) Representative images of strain analysis and quantification of (**C**) left ventricular ejection fraction (LVEF), (**D**) global longitudinal strain (GLS), (**E**) left ventricular end diastolic volume (LVEDV) and end systolic volume (LVESV). (**F**) Heart weight (HW) to tibia length (TL) ratio in ic-MI mice treated with PBS or G-CSF. (**G**) Cardiac mRNA expression of *Nppa*, *Myh6*, *Myh7* and *Il6* measured by qPCR and shown relative to baseline (BL) expression. (**H-J**) Representative images of heart section stained with Masson’s trichrome (**H**) and quantification of infarct size (**I**) and fibrosis (**J**). Data are shown as mean ± SEM. Welch’s *t*-test (**C, D**, **E**, **I** and **J**) or Mann-Whitney (**F**) or One-way ANOVA with Bonferroni *post hoc* test (**G**); * *p* < 0.05, ** *p* < 0.01 and *** *p* < 0.001. (**C-J**) BL *n* = 4, ic-MI+PBS *n* = 6-7, and ic-MI+G-CSF *n* = 5. Scale bar: 1 mm. ns, non-significant. BL, baseline mice; ic-MI, intact-chest myocardial infarction.

In line with these functional alterations, gene expression analysis in the infarcted hearts from G-CSF-treated mice revealed changes of cardiac stress markers. This was characterized by upregulation of *Nppa* (atrial natriuretic peptide, ANP), and a shift in myosin heavy chain isoform expression, with decreased *Myh6* and increased *Myh7* levels (Fig. 7G). Moreover, inflammation was sustained, as evidenced by higher *Il6* expression in hearts of G-CSF-treated animals (Fig. 7G). Histological analysis further demonstrated adverse structural remodeling, with a greater extent of fibrosis and larger scar size in hearts from G-CSF-treated mice compared with vehicle controls (Fig. 7H–J).

Overall, these findings suggest that elevating immature neutrophil numbers during the early post-MI phase leads to maladaptive remodeling with worsened cardiac performance at later stages.

### Immune landscape of the infarcted heart

Having characterized neutrophil dynamics following MI, we next investigated the responses of other leukocyte populations. Previous studies using the open-chest model have reported the presence and pathological roles of various immune cells in the infarcted heart^2-4^. However, the substantial systemic inflammation induced by the surgical procedure may profoundly influence leukocyte recruitment and activation. The intact-chest model offers the opportunity to more accurately define the immune landscape of the infarcted myocardium under conditions that minimize surgical inflammation.

Using multiparametric flow cytometry, we quantified major immune cell populations in the heart (Fig. 8A) and compared the kinetics of leukocyte infiltration between ic-MI and oc-MI models (Fig. 8B-G). We found that neutrophil infiltration peaked at levels more than three-fold higher in the oc-MI model compared to minimal-invasive ic-MI and remained elevated at day 3 post-MI (Fig. 8B). Surprisingly, classical monocytes at day 1 post-MI were more abundant in ic-MI hearts (Fig. 8C). In the later phase, macrophages increased dramatically in oc-MI hearts at days 5 and 7, whereas their numbers remained largely unchanged in ic-MI (Fig. 8D). In contrast, CD11c^+^ cells were more prominent in ic-MI hearts peaking at days 3 and 5 before declining (Fig. 8E).

**Figure 8.**
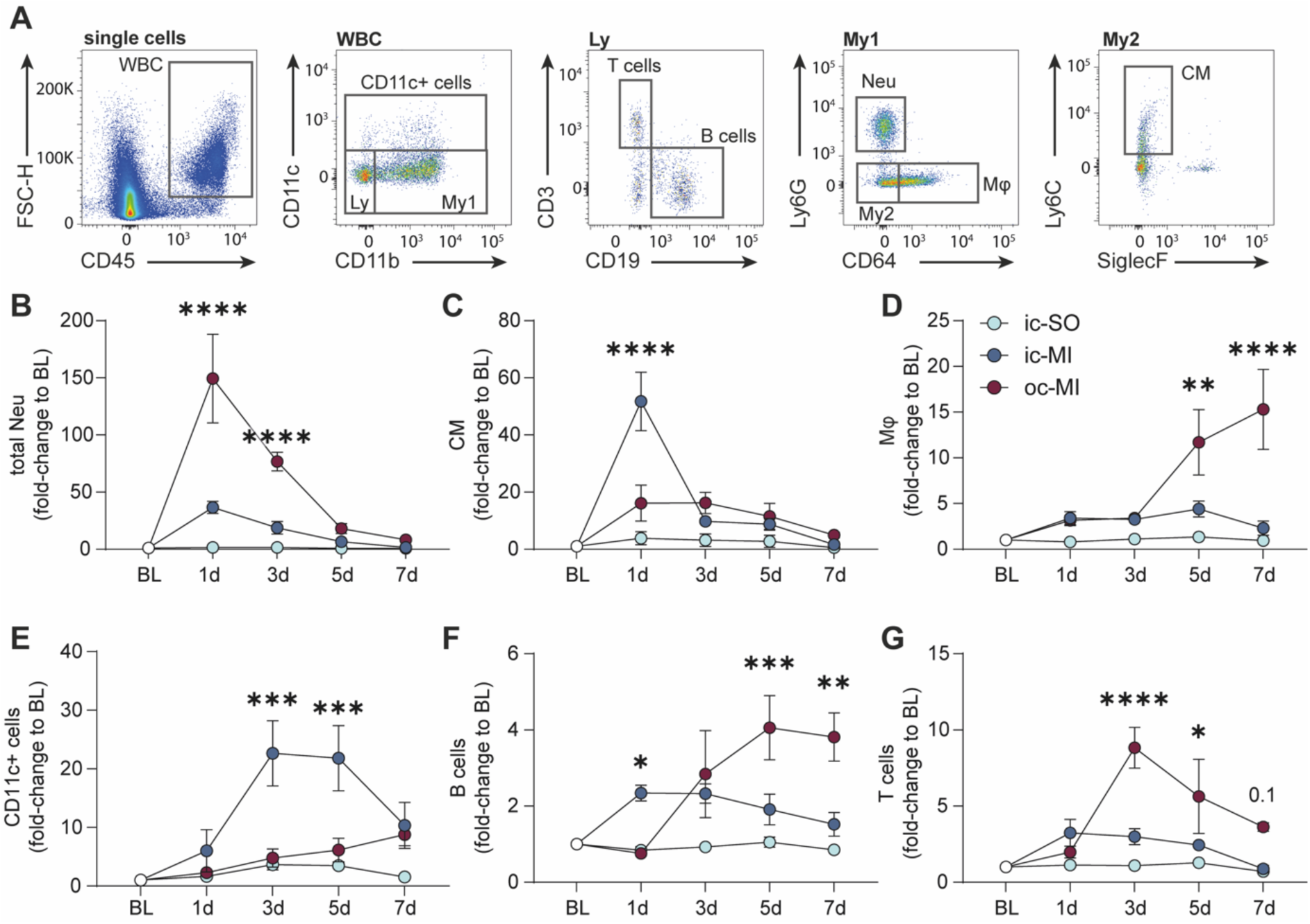
Immune landscape of the infarcted heart. Mice were subjected to myocardial infarction using either the open-chest (oc-MI, dark red) or intact-chest model (ic-MI, dark blue). Some mice were sham-operated (ic-SO, light blue). Hearts were collected at the indicated time points (**A**) Representative flow cytometry analysis to identify cardiac leukocyte subsets in heart sample following ic-MI. (**B-G**) Time courses of cardiac neutrophils (**B**), classical monocytes (**C**), macrophages (**D**), CD11c^+^ cells (**E**), B cells (**F**), and T cells (**G**). Each leukocyte population in ic-SO (light blue), ic-MI (dark blue), and oc-MI mice (red) is expressed as fold change to baseline controls (BL). Data are shown as mean ± SEM. Two-way ANOVA with Tukey’s *post hoc* test with \**p* < 0.05, \*\**p* < 0.01, \*\*\**p* < 0.001 and \*\*\*\**p* < 0.0001. *n* = 3-7 per time point per group except ic-MI 24h *n* = 10. CM, classical monocytes; MΦ, macrophages. ns, non-significant. BL, baseline mice; oc-MI, open-chest myocardial infarction; ic-SO, intact-chest sham-operation; ic-MI, intact-chest myocardial infarction.

Lymphocyte dynamics also differed between models. In minimal-invasive ic-MI hearts, B cells peaked at day 1 and returned to baseline by day 7; whereas in oc-MI hearts, B cell infiltration began at day 3, progressively increased, and reached maximal accumulation at day 5, plateauing through day 7 (Fig. 8F). We found that T cell recruitment was minimal in ic-MI hearts during the first week post-MI, while in oc-MI hearts, T cell numbers peaked at day 3 and gradually declined thereafter (Fig.8G).

Overall, these data indicate that the immune landscape differs between the intact-chest and open-chest MI models in magnitude, timing, and composition. These findings highlight the value of adopting the minimal-invasive ic-MI model for future studies of post-MI immunity.

## Discussion

Here, we provide the first comprehensive immunological validation of the minimally invasive intact-chest MI model developed by Sicklinger and colleagues^28^, and demonstrate that it faithfully recapitulates the neutrophil response observed in MI patients. We further identify immature neutrophils as contributors to post-MI outcomes. In addition, our findings highlight significant limitations of the widely used open-chest MI model for studying immune responses following MI.

The intact-chest MI model was originally developed to provide technical advantages and improve animal welfare^28^. While its superiority in terms of speed and accuracy has been well demonstrated, its impact on immune responses compared with the open-chest approach had not yet been evaluated. By directly comparing the dynamics of the neutrophil response between these two models, we found that surgical trauma profoundly reshaped neutrophil maturation in the open-chest setting. Specifically, after an initial release of nearly all mature neutrophils from BM, the open-chest procedure per se initiated hallmark features of emergency granulopoiesis. These included sustained elevations of the cytokines (G-CSF and CXCL1), a pronounced degenerative left shift with up to 80% immature neutrophils in circulation, prolonged mobilization of immature cells beyond 72 hours, and persistent alterations in the BM requiring more than a week for recovery. This is in agreement with earlier studies showing that surgical procedures in cardiovascular disease models alter myeloid cell populations^30^, including the induction of robust neutrophil mobilization^31^. It should be noted that signs of emergency granulopoiesis have been observed in previous MI studies^32^, however the contribution of the surgical procedure itself to this response had not been evaluated. Collectively, our findings reveal a major translational challenge, as surgical trauma inherent to the open-chest model induces systemic inflammation that masks disease-specific immune responses.

By contrast, we found that the minimally invasive intact-chest model provides a more pathophysiologically relevant alternative. In this setting, MI itself elicited only an acute and mild inflammatory response, characterized by transient cytokine elevation, preserved BM homeostasis, and a modest left shift that closely mirrored the neutrophil profile observed in human MI patients^23^ (Fig. 1). In this context, the MI-induced left shift most likely resulted from rapid mobilization of BM reserves rather than emergency granulopoiesis. Consequently, interventions targeting emergency granulopoiesis may be ineffective against the physiological mobilization that occurs in actual MI. This may explain the frequent failure of anti-inflammatory therapies that appear promising in preclinical studies using conventional open-chest models^8,33,34^.

To clarify post-MI neutrophil dynamics, especially immature neutrophil mobilization, we examined the spatiotemporal sequence of events following ischemia. We observed an early peak of neutrophil-mobilizing cytokines (G-CSF and IL-6) at 4 hours post-MI, followed by BM neutrophils egress at 12 hours, resulting in a marked rise in circulating immature neutrophils (≈ 32% of total neutrophils). This pattern closely aligns with previous findings from Ulrich *et al*.^17^, showing that a single G-CSF injection causes rapid mobilization of immature neutrophils, peaking at 12 hours at 28% of total neutrophils. The similarity in both timing and magnitude, between Ulrich’s study and our MI data, strongly suggests that G-CSF is the main driver of immature neutrophil release from the M after MI. Mechanistically, G-CSF is known to disrupt the CXCL12–CXCR4 retention axis in the BM, thereby promoting neutrophil egress^35^. IL-6 was also shown to mobilize immature neutrophils^36^, albeit more modestly (≈ 8% at 12 hours). Given that cytokines can act additively^37,38^, the simultaneous rise of G-CSF and IL-6 after MI likely explains the rapid, transient release of immature neutrophils. Additionally, we observed a rapid increase in plasma CXCL1 post-MI which may contribute to immature neutrophil release as CXCL1 has been shown to cooperate with G-CSF in neutrophil mobilization^39^. Together, these findings indicate that myocardial ischemia alone generates a moderate inflammatory stimulus that primarily mobilizes pre-existing bone marrow neutrophil pools, rather than inducing emergency granulopoiesis.

Once released into the bloodstream, we found that immature neutrophils migrated to the damaged myocardium as effectively as mature neutrophils with peak accumulation at 12-24 hours post-MI. Previous studies have demonstrated that immature neutrophils possess the functional migratory machinery required to infiltrate tumors^15^, we show here that they are also capable of infiltrating ischemic cardiac tissue. To date, the best-established routes of leukocyte entry into the infarcted heart include transmigration through the microvasculature in the border zone and infiltration through adipose tissue connected to the epicardium^40-42^.

Notably, a recent study using cutting-edge spatial multi-omics technologies with subcellular resolution identified a previously unrecognized route of monocyte infiltration into the infarcted myocardium via the endocardium^29^. Reanalysis of the antibody-based Sequential Immunofluorescence dataset from this study revealed that neutrophils were localized not only to the border zone and the region adjacent to the epicardium, but also beneath the endocardial surface. This raises the possibility that neutrophils, like monocytes, may be recruited through the endocardium. Further investigations will be required to delineate the molecular pathways governing the distinct routes of neutrophil trafficking.

To specifically assess the functions of immature neutrophils post-MI, we enhanced their mobilization using G-CSF/anti–G-CSF antibody complexes, which prolong G-CSF half-life and enhance its biological activity^15,43^. We showed that, after MI, G-CSF treatment increased immature neutrophil numbers in blood and proportionally in infarcted tissue. This accumulation was associated with impaired cardiac function, larger infarcts, and pathological remodeling. These results establish a causal role for immature neutrophils in maladaptive remodeling. Importantly, this has direct clinical relevance as a recent study showed that elevated circulating immature granulocytes in MI patients is associated with a more than 5–6-fold increase in 30-day mortality risk^24^. Together, these findings highlight that immature neutrophils are not mere bystanders but rather act as key mediators of post-infarct pathology and may serve as valuable biomarkers for risk stratification and outcome prediction.

Neutrophils contribute to tissue repair by clearing dead cells, but their effector functions can also cause collateral damage. A growing body of evidence suggests that immature neutrophils play pathological roles in cancer^13,44^ and their accumulation in tumors correlates with a poor prognosis^15,45-47^. By contrast, far less is known about the role of immature neutrophils in non-malignant diseases. While a previous study showed that resting immature neutrophils in the BM had limited effector capacity^15^, we demonstrated that immature neutrophils infiltrating the infarcted myocardium are functionally competent cells. Indeed, cardiac immature neutrophils mounted a robust oxidative burst, comparable to mature neutrophils and displayed phagocytic capacity. We propose here two non-mutually exclusive explanations for the observed pathological consequences on the infarcted heart associated with immature neutrophil accumulation. First, and most importantly, we would expect a higher number of competent neutrophils, regardless of maturation state, to exarcebate the overall inflammatory response. An excess of ROS-producing cells should increase oxidative stress on cardiomyocytes, endothelial cells, and extracellular matrix, thereby amplifying tissue injury^48,49^. Second, although marginal, we observed a reduced phagocytic capacity in immature neutrophils compared to mature neutrophils, which may limit to some extent the efficiency of efferocytosis, a process essential for clearing apoptotic cells and resolving inflammation. Inadequate debris clearance promotes the persistence of pro-inflammatory signals^50^, consistent with the elevated IL-6 levels we observed at 28 days post-MI. Notably, a comparable neutrophil phenotype has been described in the inflammatory vascular disease giant cell arteritis^51^, where immature neutrophils were similarly shown to produce high levels of ROS, comparable to mature neutrophils, while exhibiting a modestly reduced phagocytic capacity, and where their accumulation was associated with increased tissue damage and vascular pathology. Taken together, we propose that the higher number of total ROS-competent neutrophils (comprised from both mature and immature neutrophils) in the infarcted heart, combined with suboptimal clearance by immature neutrophils, sustains local inflammation, exacerbates tissue injury, and ultimately impairs cardiac healing and function.

Beyond neutrophils, the intact-chest MI model revealed distinct patterns of immune cell recruitment that differ in magnitude, kinetics, and composition compared with the open-chest model, suggesting that previous studies may have mischaracterized some aspects of the inflammatory response following MI^2-4,52^. For instance, we found that the accumulation of T cells, B cells, and neutrophils in the heart was markedly higher in the open-chest model, indicating that surgical trauma in the open-chest model induces a strong ischemia-independent inflammatory response. We also observed divergent profiles of monocytes, macrophages, and CD11c^+^ cells between the two models. Further work will be required to delineate the molecular pathways underlying these model-specific differences and to define the precise roles of individual immune-cell subsets in the intact-chest MI model.

In conclusion, our study underscores the importance of using pathophysiologically accurate experimental models. As George E. P. Box famously noted: "All models are wrong, but some are useful”^53^. The careful selection of models and controls has become increasingly recognized as essential for correctly defining the cellular and molecular pathways involved in cardiovascular disease^54-56^. Building upon the technical foundation established by Sicklinger and colleagues^28^, our findings demonstrate that the intact-chest MI model represents the more useful choice with respect to immune response characterization as it avoids the confounding effects of surgical trauma. Using neutrophils as a benchmark, we showed that the murine intact-chest MI model faithfully recapitulates human neutrophil responses in AMI patients and revealed the pathogenic contribution of immature neutrophils to heart failure. Although our study focused primarily on neutrophils, our findings also suggest that the ic-MI model may better reflect aspects of the broader post-MI immune landscape. We anticipate that future studies using the intact-chest MI model will clarify the roles of the different immune cell types in disease progression with higher clinical relevance and facilitate the identification of therapeutic targets specifically addressing the pathophysiological inflammatory processes of MI.

## Methods

### Ethics declarations

All experiments involving human specimens were approved by the research ethics committees of the National Healthcare Group Domain Specific Review Board (NHG DSRB Domain C 2020/01253). All procedures involving animals were performed conform to the guidelines from Directive 2010/63/EU, the government of Upper Bavaria ((ROB-55.2.2532.Vet_02-13-176, 55.2.2532.Vet_02-18-114 and 55.2.2532.Vet_23-157), Germany, and the Institutional Animal Care and USA Committee (IACUC), in accordance with the guidelines of the Agri-Food and Veterinary Authority (AVA) and the National Advisory Committee for Laboratory Animal Research (NACLAR) of Singapore. All procedures had local ethical approval, and all experiments were conducted according to the 2013 Declaration of Helsinki.

### Human data and study design

Among twenty patients with acute myocardial infarction (AMI), 13 met the inclusion criteria and were categorized into non-ST-elevation MI (NSTEMI) and ST-elevation MI (STEMI). All details regarding the patients are provided in **Supplementary Table 1**. Five healthy donors (HD) were recruited as control subjects.

### Flow cytometry of human blood samples

Venous blood was collected in EDTA tubes. For multiparameter flow cytometry, 2mL of whole blood were lysed using 20 mL of 1X RBC lysis buffer (eBioscience) for 6 min at room temperature. The lysis reaction was stopped by adding 30 mL of PBS, and cells were centrifuged at 400 g for 5 min. RBC lysis step was repeated once. Subsequently cells were blocked with anti-human CD32 antibody for 15 min, followed by fluorochrome-conjugated antibodies (**Supplementary Table 2**) for 30 min at 4 °C. Cells were washed, resuspended in PBS containing DAPI, and data was acquired on a LSRII flow cytometer (BD Biosciences). Analysis of immune cell populations and applied gating strategies were conducted with FlowJo software (v10.8.0, TreeStar). Neutrophils were defined as CD45^+^ Lin^-^ CD15^+^ CD66b^+^ CD49d^-^ CD11b^+^. CD16 and CD10 expression was used to distinguish mature (CD16^+^ CD10⁺) and immature (CD16^+/-^ CD10⁻) neutrophils. Representative gating strategy is shown in Extended Data Figure 1.

### Mouse experiments

Female C57BL/6J mice (9-12 weeks old; Janvier Labs) were housed under standard laboratory conditions with a 12-hour light/dark cycle and provided chow and water ad libitum. Animals were subjected to either sham-operation (SO) or myocardial infarction (MI). Healthy untouched mice were used as baseline controls (BL). All surgeries were performed between ZT3 and ZT5 to prevent circadian rhythm influencing neutrophil dynamics on time of surgery.

### Open-chest MI model

In the open-chest (oc) model, MI was induced via invasive surgery and permanent ligation of the left anterior descending coronary artery (LAD) as described previously^9,57^. In brief, mice were anesthetized with midazolam (5 mg/kg), medetomidine (0.5 mg/kg), and fentanyl (0.05 mg/kg), followed by endotracheal intubation and ventilation using a MiniVent ventilator (Harvard Apparatus, Holliston, MA). Thoracotomy was performed on the left side at the third intercostal space, the pericardium was opened, and the LAD was permanently ligated proximal to its bifurcation using 7-0 permahand silk threads (Ethicon, Somerville, USA). The intercostal space and skin were closed with 5-0 Seraflex silk (Serag Wiessner, Germany). Anesthesia was reversed with flumazenil (0.5 mg/kg), and atipamezole (2.5 mg/kg). Analgesia (buprenorphine, 0.05 mg/kg) was administered 30 min pre as well as 6 and 18 h post-surgery. Sham-operated (oc-SO) animals underwent the same procedure without LAD ligation.

### Intact-chest MI model

In the intact-chest (ic) model, minimally invasive MI induction was performed under echocardiographic guidance as described by Sicklinger et al^28^. In brief, mice were anesthetized with isoflurane (1 L O₂/min; 3-4% for induction and 1-2% for procedure) and positioned on a Vevo imaging station (Vevo3100, VisualSonics Inc,). The LAD coronary artery was visualized at the mid-papillary level using Color Doppler imaging. A monopolar needle attached to a micromanipulator was introduced through the chest and placed directly onto the LAD. Artery vessel was coagulated with high-frequency electricity (HF-Gerät Erbe ICC 200). Successful MI induction was confirmed by persistent loss of Doppler flow and akinesia of the affected left ventricular region. Analgesia (buprenorphine, 0.1 mg/kg) was administered 30 min pre and 6 h post-surgery. Sham-operated (ic-SO) animals underwent the same procedure without electrical coagulation.

### Organ collection and processing

Animals were euthanized with ketamine/xylazine overdose (*i.p*.; 150/30 mg/kg) at respective time points post-MI (4h, 12h, 24h, 72h, 5d, 7d). Whole blood was collected via cardiac puncture in EDTA-coated tubes. After transcardiac perfusion with ice-cold PBS to remove remaining blood within the heart and vessels, the following organs: tibia, femur, spleen, mediastinal lymph nodes (MLNs), peripheral lymph nodes (axillary and inguinal; LN), left lung lobe and hearts (without atria) were excised and further processed. Tibia length was measured to normalize heart weight.

### Flow cytometry

Cardiac tissues were minced with fine scissors and digested with Collagenase Type I (450 U/mL), Collagenase Type XI (125 U/mL), DNase I (60 U/mL) and Hyaluronidase (60 U/mL) for 1h at 37°C. Bone marrow (BM) cells were isolated from one femur by removing the epiphysis and centrifuging for 2 min at 10’000 rpm. Lungs were minced with fine scissors and digested with Collagenase Type I (450 U/mL), Hyaluronidase (120 U/mL), and DNase I (240 U/mL) for 45min at 37 °C. Spleens, MLN and peripheral LNs were mashed through a 50 µm filter. Red blood cells in blood, spleen and lung were lysed using RBC lysis buffer (BioLegend). All cells were washed and resuspended in FACS Buffer (PBS containing 1% BSA). Fc receptors were blocked with rat anti-mouse CD16/32 antibody (clone 93, BioLegend) and surface proteins stained for 45 min at 4°C with antibodies (**Supplementary Table 2).** Following washing, cells were resuspended in FACS Buffer containing CountBright Absolute Counting Beads (Life Technologies) and measured with an LSRFortessa X-20 flow cytometer (BD Biosciences). Analysis of immune cell populations and applied gating strategies were conducted with FlowJo software (v10.8.0, TreeStar). Neutrophils were defined as CD45^+^ CD11b^+^ CD115^-^ Ly6G^+^ in blood, Lineage^-^ CD45^+^ CD11b^+^ CD115^-^ Ly6G^+^ in bone marrow, and CD45^+^ CD11b^+^ Ly6G^+^ in the heart. CD101 expression was used to distinguish mature (CD101⁺) and immature (CD101⁻) neutrophils. A CD101 fluorescence minus one (FMO) control was included to accurately define positive and negative populations. Representative gating strategies and FMO stainings are shown in **Extended Data Figures 2 and 5**. CXCR2 expression in BM immature neutrophils was quantified as geometric mean fluorescence intensity (MFI). For comparative analysis, MFI values from BM cells of oc-SO, oc-MI, ic-SO, and ic-MI groups were normalized to the MFI of the respective marker in immature neutrophils from the baseline (BL) group.

### ROS production and phagocytosis assay

Reactive oxygen species (ROS) production was quantified by incubating cells with 5 µM Dihydrorhodamine 123 (DHR 123, Sigma-Aldrich) for 10 min at 37°C. Oxidation of DHR to fluorescent rhodamine 123 was measured by flow cytometry (FITC channel). Phagocytic activity was assessed using pHrodo™ Deep Red *E. coli* BioParticles™ (Invitrogen). Cells were incubated with 40 µg of opsonized BioParticles for 30 min at 37°C. Upon internalization into acidic phagosomes, pHrodo fluorescence increases, allowing quantification of phagocytosis by flow cytometry (Deep Red channel). Following incubation, cells were washed with FACS buffer and stained with fluorochrome-conjugated antibodies as described above. After a final wash, cells were resuspended in FACS buffer and acquired on an LSRFortessa X-20 flow cytometer (BD Biosciences). Data were analyzed using FlowJo software (v10.8.0, TreeStar).

### Increased immature neutrophils recruitment via G-CSF injection

To increase circulating immature neutrophil counts, mice were treated with G-CSF/anti–G-CSF antibody complexes, which prolong G-CSF half-life and enhance its biological activity, as previously described by Rubinstein et al.^43^ and Evrard et al.^15^. Briefly, human G-CSF (PeproTech) was incubated with rat anti-human G-CSF antibody (clone BVD11-37G10, Southern Biotech) at a 1:6 ratio for 20 min at 37 °C. The resulting complex was diluted in sterile PBS to a final dose of 1.5 µg G-CSF in 100 µL per mouse. Mice received a single intraperitoneal injection of the complex or vehicle (PBS) immediately after MI induction in the intact-chest model. Tissues were harvested 24h or 28 days post-MI, as described above.

### Cytokine and chemokine immunoassays

Plasma was collected by centrifuging blood for 10 min at 10’000 rpm. Cytokine and chemokine levels were assessed using a custom-made ProcartaPlex (ThermoFisher) containing GROα/CXCL1, MCP-1/CCL2, G-CSF, IL-6, and TNFα, as well as DuoSet ELISAs for mouse CXCL1, G-CSF, and IL-6 (R&D Systems) following manufacturers’ protocols.

### Gene expression analysis

Cardiac apex and base of G-CSF and PBS injected mice (28d post-MI) were snap frozen. Frozen hearts were first pestilled in liquid nitrogen to a fine powder, then lysed in RLT Plus buffer (Qiagen) supplemented with β-mercaptoethanol and homogenized using a bead mill (Ø7mm). RNA was isolated with RNeasy Plus Micro Kit (Qiagen) following manufacturer’s protocol. Using the PrimeScript RT Reagent Kit (TaKaRa Bio), 1 µg of RNA was reverse transcribed and 25 ng of the resulting cDNA per reaction was used for TaqMan real-time RT-PCR analysis. mRNA expression was determined for *Nppa* (Mm01255747_g1), *Myh6* (Mm00440359_m1)*, Myh7* (Mm00600555_m1) and *Il6* (Mm00446190_m1) (Applied Biosystems). Samples were run in duplicates using a QuantStudio 6 Pro Real-Time-PCR-System (ThermoFisher) and mRNA expression was normalized to *Gapdh* levels (Mm99999915_g1, Applied Biosystems). Relative gene expression was calculated with the comparative ΔΔCT method.

### Histology

Mid-section of hearts from G-CSF and PBS injected mice (28d post-MI) were embedded in Tissue-Tek Optimal Cutting Temperature (O.C.T) compound in a plastic cassette and were immediately snap frozen. Hearts were stored at -80 °C and further sectioned at 5μm thickness using a Leica CM3050S cryostat. Tissue sections were stained with a Masson’s Trichrome (Sigma-Aldrich) according to the manufacturer’s instructions. Brightfield images were acquired using a Leica Thunder microscope (Leica Microsystems) and analyzed with Fiji/ImageJ (https://imagej.net). For each data point, three sections from the apex, mid, and proximal infarct area were analyzed and averaged. Scar size measurement: scarred regions were manually delineated based on blue collagen staining, measured, and normalized to the total left ventricular area. Fibrosis content measurement: fibrosis was quantified as the proportion of collagen-positive tissue relative to the total left ventricular area. The left ventricle was manually outlined to define the total ventricular area, and blue-stained collagen was isolated using color thresholding (Chanel_1 in Colour Deconvolution plugin).

### Echocardiography

Echocardiographic scans of the parasternal long axis were obtained 28 days post-MI from mice treated with G-CSF or PBS using the Vevo 3100 imaging system (VisualSonics Inc.). Mice were anesthetized with isoflurane (3-4% for induction and 1-2% for acquiring scans) at a flow rate of 1L O₂/min. Heart rate and body temperature were kept stable throughout the procedure. Left ventricular end-diastolic volume, end-systolic volume and ejection fraction were measured based on the left parasternal long-axis view using VevoLab software version 5.10.0 (VisualSonics). Global longitudinal strain was quantified in the longitudinal axis by speckle tracking using VevoStrain.

### Mapping of neutrophils in cardiac tissue

To assess the spatial distribution of neutrophils within infarcted cardiac tissue, we analyzed data from the open-source SeqIF dataset generated by Wünnemann et al.^29^. In that study, cryosections (8 μm) of mouse hearts collected 24 h after intact-chest MI were stained with a 12-antibody panel using the Lunaphore COMET platform (Lunaphore Technologies). We downloaded the raw TIFF files from Synapse (https://www.synapse.org, accession: syn51449058) and high-resolution images (30,238 × 28,381 pixels) containing MPO, Tnnt2, and Ankrd1 staining were selected and processed using Fiji/ImageJ.

### Statistical Analysis

After performing a normality test, an unpaired Welch’s *t*-test or Mann-Whitney test was performed for comparisons between two groups. For comparisons within three groups, a one-way ANOVA followed by Bonferroni’s *post hoc* was performed. For comparisons within two or more groups within different time points, a two-way ANOVA with Šidák’s or Tukey’s *post hoc* if appropriate was used. Paired or unpaired multiple *t*-tests with Holm–Šidák’s correction were used for comparisons between ic-SO and ic-MI groups. All details of the statistical tests used are provided in the figure legends. Statistical significance is represented by asterisks and the corresponding *p* values, as indicated in the legends. Data are expressed as mean ± SEM. Biological replicates are stated in the legends for each figure. All statistical analysis were performed with Prism10 (v10.4.1, GraphPad). Heat maps were created with Morpheus analysis software (https://software.broadinstitute.org/morpheus).

## Acknowledgments

We acknowledge the support by Deutsche Forschungsgemeinschaft (DFG) (SFB1123-A1/A10 to J.D. and C.W., SFB1123-B09, STE1053/5-1, STE1053/8-1 to S.S.), by the German Center for Cardiovascular Research (DZHK) (81X2600271, 81X2600275 and 81X2600283 to J.D., 1X2600254 to C.W. and 81Z0600205 to S.S.), by Förderprogramm für Forschung und Lehre (FöFoLe) (68/2013 to S.S). I.K. was supported by the Singapore Immunology Network (SIgN) Agency for Science, Technology and Research (A*STAR).

## Author contributions

Z.M.R. and A.K. designed experiments, performed mouse experiments, flow cytometry experiments and echocardiography measurements, analyzed data, and contributed to writing the manuscript. S.L.P. and M.S. performed mouse experiments and analyzed data. Y.J. performed tissue embedding/sectioning. A.E. performed gene expression experiments. K.N. performed flow cytometry experiments and analyzed data. I.K. and M.Y.Y.C. provided and analyzed data from human AMI samples and provided intellectual input. F.S. and F.L. provided assistance with intact-chest mouse model and provided intellectual input. C.W. provided intellectual input and funding. S.S. provided strategic input and funding, obtained animal experimentation approvals and supervised research enabling the implementation of MI mouse models. J.D. designed and supervised the study, planned experiments, provided funding, and wrote the manuscript with input from all authors.

