## Supplemental Information for "Minimally invasive myocardial infarction model recapitulates patient immune responses and reveals a pathogenic role for immature neutrophils"

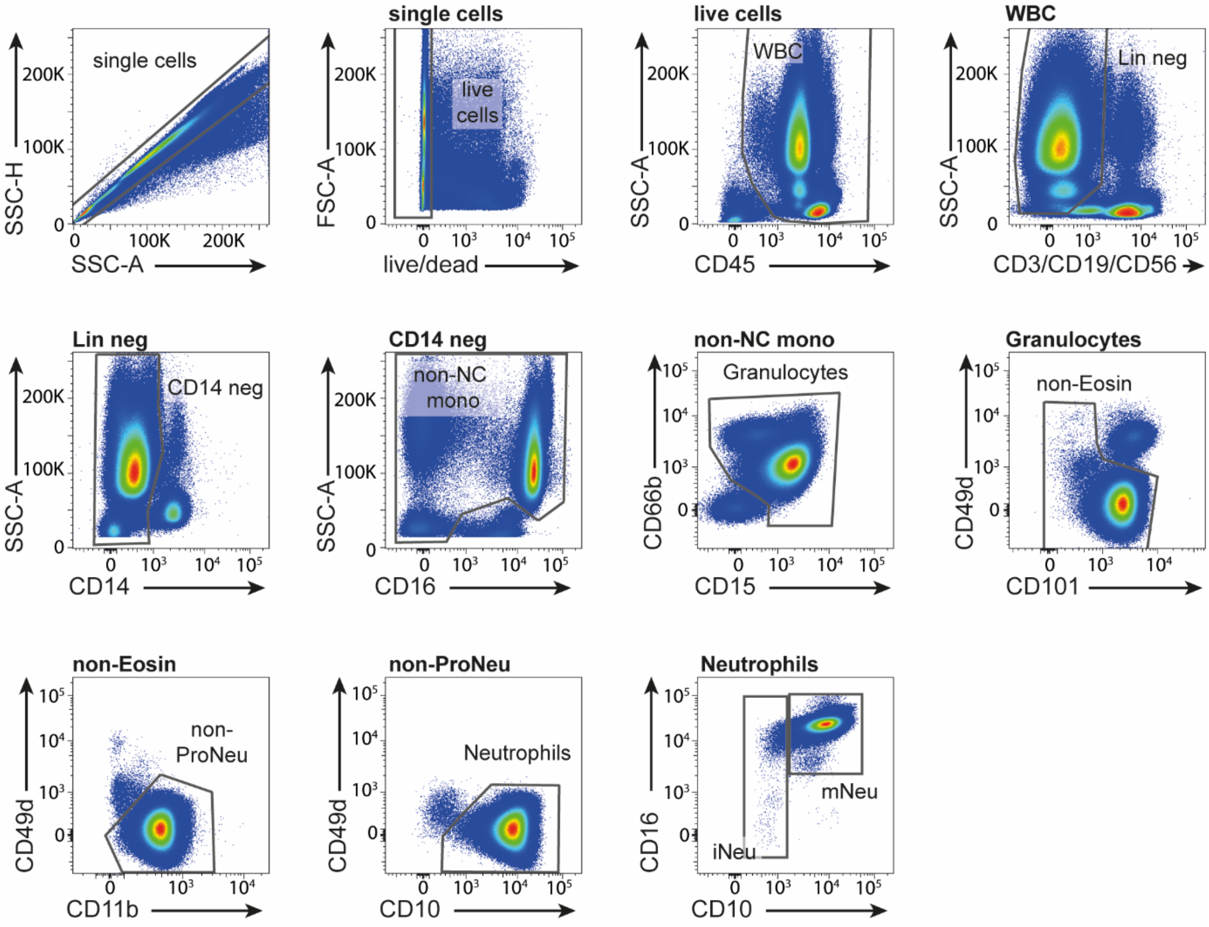


**Extended Data Figure 1. Flow cytometry gating strategy for human neutrophil subsets.** Flow cytometry gating strategy to identify circulating iNeu and mNeu subsets in AMI patients. Neutrophils are defined as CD45$^{+}$, lineage (CD3/CD19/CD56)$^{-}$, CD14$^{-}$, SSC$^{high}$, CD15$^{+}$, CD66b$^{+}$, CD101$^{+}$, and CD49d$^{-}$. iNeus are identified as CD16$^{-/+}$ CD10$^{-}$ and mNeus as CD16$^{+}$ CD10$^{+}$. AMI, acute myocardial infarction; mNeu, mature neutrophils; iNeu, immature neutrophils.

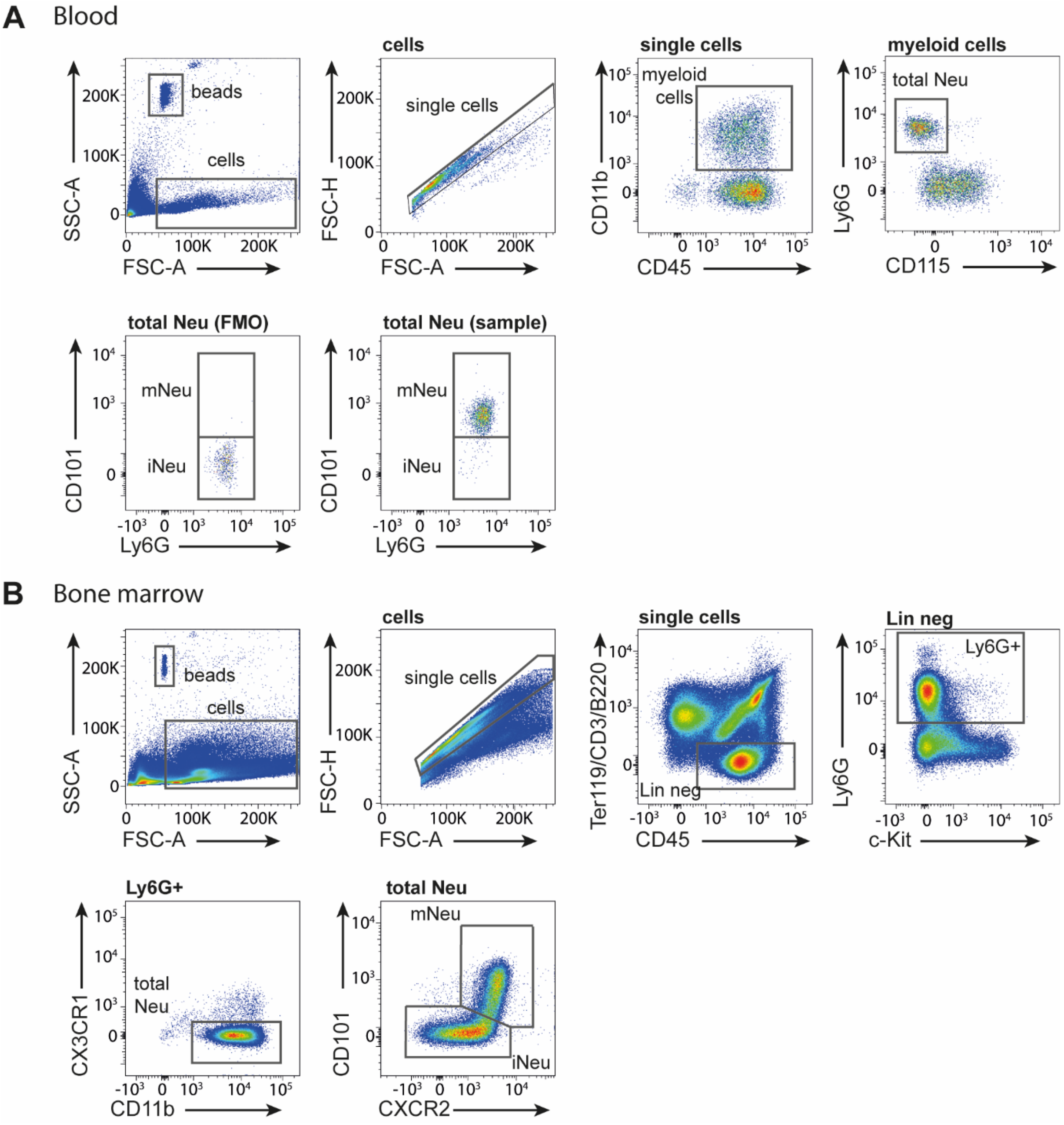


**Extended Data Figure. 2. Flow cytometry gating strategy for murine leukocyte subsets.**

Flow cytometry gating strategy to identify neutrophil subsets in murine tissues. (**A**) Blood neutrophil (CD45$^+$CD11b$^+$Ly6G$^+$) subsets were defined by CD101 expression based on FMO. (**B**) Bone marrow neutrophil (CD45$^+$Lin$^-$Ly6G$^+$c-Kit$^{-/+}$CX3CR1$^-$CD11b$^+$) subsets were defined by CD101 and CXCR2 expression. WBC, white blood cells; Neu, neutrophils; iNeu, immature neutrophils; mNeu, mature neutrophils.

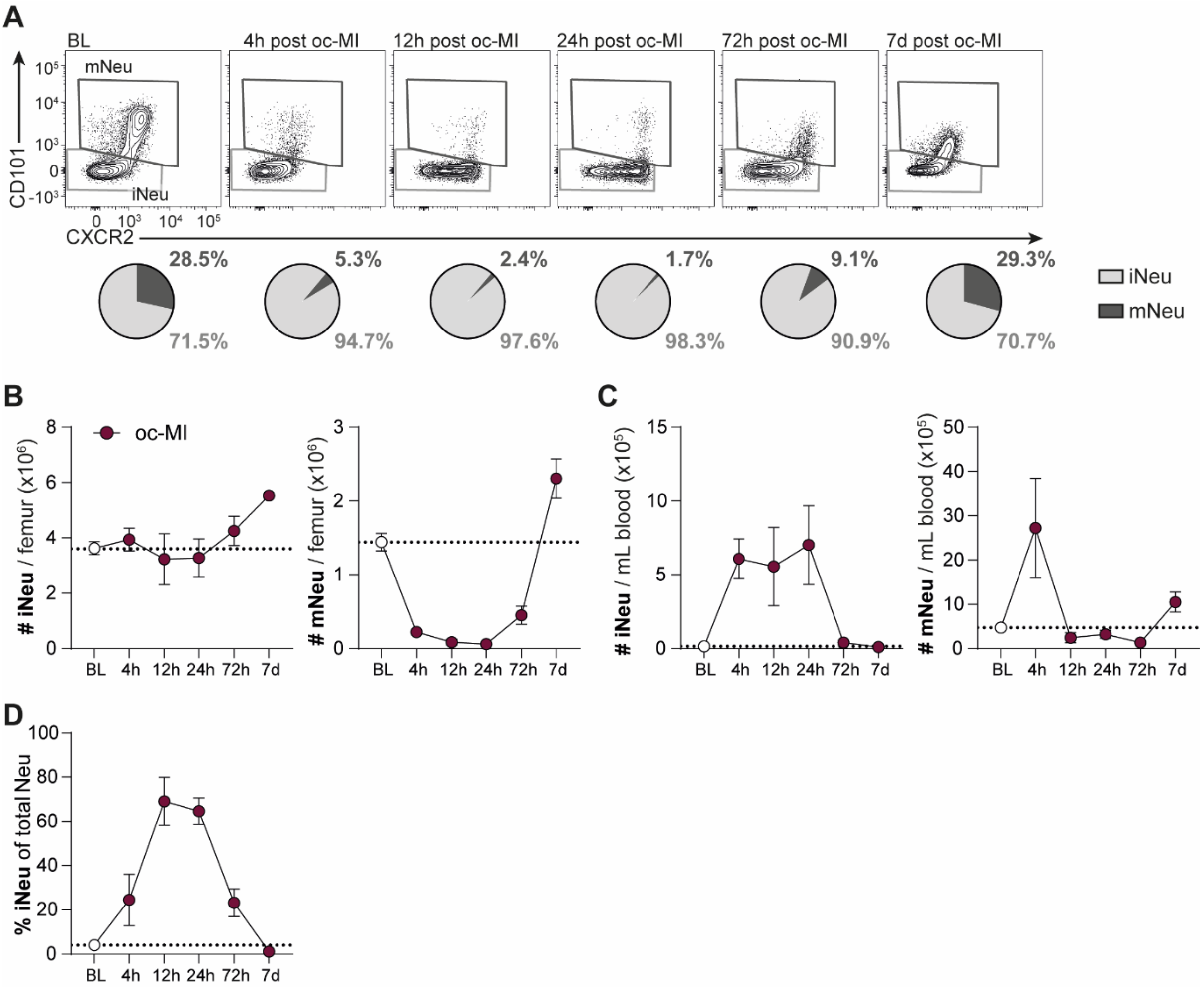


**Extended Data Figure. 3. Time-course of neutrophil subsets in oc-MI model.**

(**A**) Representative flow cytometry plots of neutrophil subsets in bone marrow (BM) of BL mice and 4h-7d post oc-MI. Pie charts represent average percentages of subsets within total BM neutrophils. (**B**) Absolute counts of BM neutrophil subsets at different time points after oc-MI. (**C**) Absolute counts of circulating neutrophil subsets at different time points after oc-MI. (**D**) Circulating iNeu fraction within total neutrophil population at different time points post oc-MI. Data are shown as mean ± SEM. Multiple *t*-test between BL and oc-MI groups for each time-point corrected with Holm-Šidák *post hoc* test; *$p < 0.05$, **$p < 0.01$ and ***$p < 0.001$. All time points in BM and blood contain $n = 3$ per group except BL $n = 15$. BL, baseline mice; oc-MI, open-chest myocardial infarction; mNeu, mature neutrophils; iNeu, immature neutrophils.

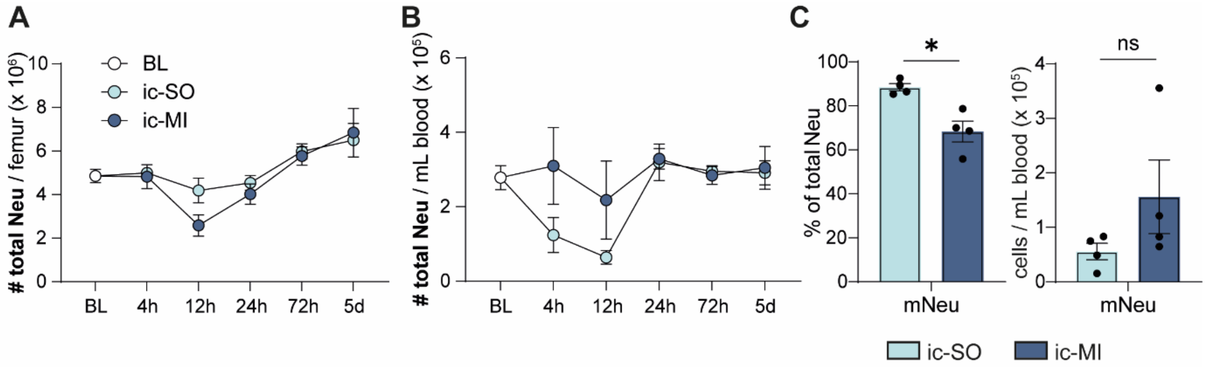


**Extended Data Figure. 4. Dynamics of neutrophil subsets in bone marrow and blood in ic-MI model.**

(**A**) Time course of bone marrow (BM) neutrophil counts in ic-SO and ic-MI mice. (**B**) Time course of circulating neutrophil counts in ic-SO and ic-MI mice. (**C**) Fraction of total neutrophils (left) and counts (right) of mature neutrophils in blood 12h post-intervention. Data are shown as mean ± SEM. (**A** and **B**) Two-way ANOVA with Šidák *post hoc* test; $n$ = 3-6 per time point per group, except ic-SO 24h $n$ = 7 and ic-MI 24h $n$ = 11. (**C**) Welch's $t$-test or Mann-Whitney test; *$p < 0.05$. ic-SO $n$ = 4 and ic-MI $n$ = 4. ns, non-significant, mNeu, mature neutrophils; ic-SO, intact-chest sham-operation; ic-MI, intact-chest myocardial infarction.

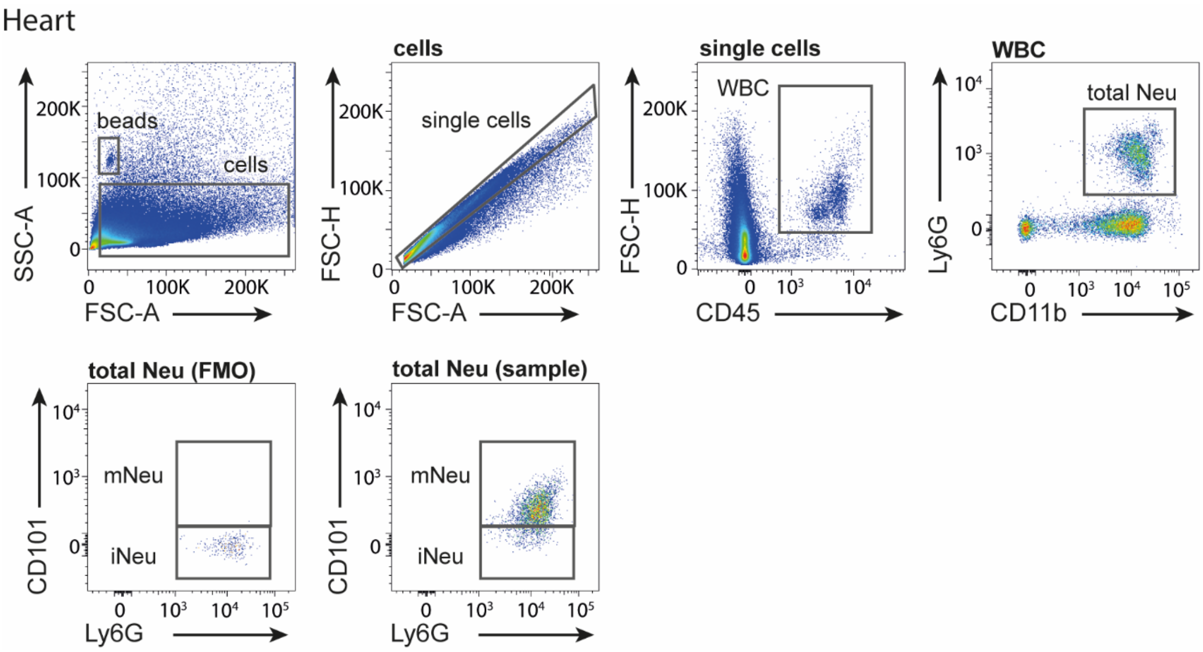


**Extended Data Figure 5. Flow cytometry gating strategy for cardiac leukocyte subsets.** Flow cytometry gating strategy to identify neutrophil subsets in murine hearts gated on ic-MI sample. Cardiac neutrophil ($CD45^{+}CD11b^{+}Ly6G^{+}$) subsets were defined by CD101 expression based on FMO. WBC, white blood cells; Neu, neutrophil; iNeu, immature neutrophil; mNeu, mature neutrophil; Ly6C$^{hi}$, classical Ly6C$^{hi}$ monocytes in BM; CM, classical Ly6C$^{hi}$ monocytes.

| | | AMI<br>(*n* = 20) | NSTEMI<br>(*n* = 10) | STEMI<br>(*n* = 10) |
|---|---|---|---|---|
| **Age (years)** | | 57.7 ± 9.5 | 60.2 ± 9.9 | 55.2 ± 9.0 |
| **Gender** | Male/Female | 16/4 | 8/2 | 8/2 |
| **BMI (kg/m2)** | | 26.5 ± 4.1 | 27.9 ± 4.3 | 25.1 ± 3.6 |
| **Analysis** | LVEF (%) | 45.4 ± 13.1 | 43.3 ± 14.0 | 47.4 ± 12.5 |
| | Troponin (ng/L) | 11070.7 ± 9757.9 | 6967.1 ± 9552.7 | 15174.2 ± 9510.1 |

**Supplementary Table 1. General traits from AMI patients.**

Data are represented as mean ± SD. AMI, acute myocardial infarction; NSTEMI, non-ST elevated myocardial infarction; STEMI, ST elevated myocardial infarction; LVEF, left ventricular ejection fraction.

**Supplementary Table 2. List of antibodies.**

| Reagent or Resource | Fluorochrome | Clone | Source | Cat. No |
|---|---|---|---|---|
| Anti-human CXCR2 | FITC | 5E8/CXCR2 | BioLegend | 320704 |
| Anti-human CD10 | PE | HI10a | BioLegend | 312203 |
| Anti-human CD49d | PE Dazzle 594 | 9F10 | BD Biosciences | 563645 |
| Anti-human CD3 | PE-Cy7 | UCHT1 | BioLegend | 300406 |
| Anti-human CD19 | PE-Cy7 | HIB19 | BioLegend | 302206 |
| Anti-human CD56 | PE-Cy7 | HCD56 | BioLegend | 318304 |
| Anti-human HLA-DR | BV605 | L243 | BioLegend | 307640 |
| Anti-human CD15 | BV650 | HI98 | BD Biosciences | 564232 |
| Anti-human CD14 | BV711 | M5E2 | BioLegend | 301838 |
| Anti-human CD45 | APC-Cy7 | HI30 | BioLegend | 304014 |
| Anti-human CD11b | BUV395 | ICRF44 | BD Biosciences | 563839 |
| Anti-human CD16 | BUV737 | 3G8 | BD Biosciences | 612786 |
| Anti-mouse B220 | PE-Cy5 | RA3-6B2 | BioLegend | 103210 |
| Anti-mouse CD3 | BUV737 | 17A2 | BD Biosciences | 612803 |
| Anti-mouse CD3 | PE-Cy5 | 17A2 | BioLegend | 100274 |
| Anti-mouse CD11b | APC-Cy7 | M1/70 | BioLegend | 101226 |
| Anti-mouse CD11b | PE Dazzle 594 | M1/70 | BioLegend | 101256 |
| Anti-mouse CD11c | BV510 | N418 | BioLegend | 117353 |
| Anti-mouse CD11c | BV786 | N418 | BioLegend | 117336 |
| Anti-mouse CD16/32 | unconjugated | 93 | BioLegend | 101302 |
| Anti-mouse CD16/32 | APC-Cy7 | 93 | BioLegend | 101328 |
| Anti-mouse CD19 | PE Dazzle 594 | 6D5 | BioLegend | 115554 |
| Anti-mouse CD45 | BV605 | 30-F11 | BioLegend | 103155 |

| | | | | |
|---|---|---|---|---|
| Anti-mouse CD45 | BV711 | 30-F11 | BioLegend | 103147 |
| Anti-mouse CD64 | BV421 | X-54-5/7.1 | BioLegend | 139309 |
| Anti-mouse CD64 | BV711 | X-54-5/7.1 | BioLegend | 139311 |
| Anti-mouse CD62L | BV786 | MEL-14 | BioLegend | 104440 |
| Anti-mouse CD101 | PE | Moushi101 | Invitrogen | 12-1011-82 |
| Anti-mouse CD101 | PE-Cy7 | Moushi101 | Invitrogen | 25-1001-82 |
| Anti-mouse CD115 | APC | AFS98 | BioLegend | 135510 |
| Anti-mouse CD115 | BUV395 | T38-320 | BD Biosciences | 743642 |
| Anti-mouse c-Kit | BV786 | ACK2 | BioLegend | 135138 |
| Anti-mouse c-Kit | PE-Cy7 | 2B8 | BioLegend | 105814 |
| Anti-mouse CXCR2 | BUV737 | V48-2310 | BD Biosciences | 748680 |
| Anti-mouse CX3CR1 | BV650 | SA011F11 | BioLegend | 149007 |
| Anti-mouse Ly6C | BV510 | HK1.4 | BioLegend | 128033 |
| Anti-mouse Ly6C | BV605 | HK1.4 | BioLegend | 128035 |
| Anti-mouse Ly6C | A700 | HK1.4 | BioLegend | 128024 |
| Anti-mouse Ly6C | APC-Cy7 | HK1.4 | BioLegend | 128026 |
| Anti-mouse Ly6G | BV510 | 1A8 | BioLegend | 127633 |
| Anti-mouse Ly6G | BV786 | 1A8 | BioLegend | 127645 |
| Anti-mouse Ly6G | APC-Cy7 | 1A8 | BioLegend | 127624 |
| Anti-mouse Ly6G | BUV395 | 1A8 | BD Biosciences | 565964 |
| Anti-mouse SiglecF | BV421 | E50-2440 | BD Biosciences | 565934 |
| Anti-mouse Ter119 | PE-Cy7 | TER-119 | BioLegend | 116210 |
